# Zur and Zinc Increase Expression of *E. coli* Ribosomal Protein L31 Through RNA-Mediated Repression of the Repressor L31p

**DOI:** 10.1101/2022.05.27.493739

**Authors:** Rebecca A. Rasmussen, Jeannie M. Camarillo, Victoria Sosnowski, Byoung-Kyu Cho, Young Ah Goo, Julius B. Lucks, Thomas V. O’Halloran

## Abstract

Bacteria can adapt in response to numerous stress conditions. One such stress condition is zinc depletion. The zinc-sensing transcription factor Zur regulates the way enteric bacteria respond to severe changes in zinc availability. Under zinc sufficient conditions, Zn-loaded Zur (Zn_2_-Zur) is well-known to repress transcription of genes encoding zinc uptake transporters and paralogues of a few ribosomal subunits. Here, we report the discovery and mechanistic basis for the ability of Zur to up-regulate expression of the ribosomal protein L31 in response to zinc in *E. coli*. Through genetic mutations and reporter gene assays, we find that Zur achieves the up-regulation of L31 through a double repression cascade by which Zur first represses the transcription of L31p, a zinc-lacking paralogue of L31, which in turn represses the translation of L31. Mutational analyses show that translational repression by L31p requires an RNA hairpin structure within the *l31* mRNA and involves the N-terminus of the L31p protein. This work uncovers a new genetic network that allows bacteria to respond to host-induced nutrient limiting conditions through a sophisticated ribosomal protein switching mechanism.

**Graphical Abstract:** 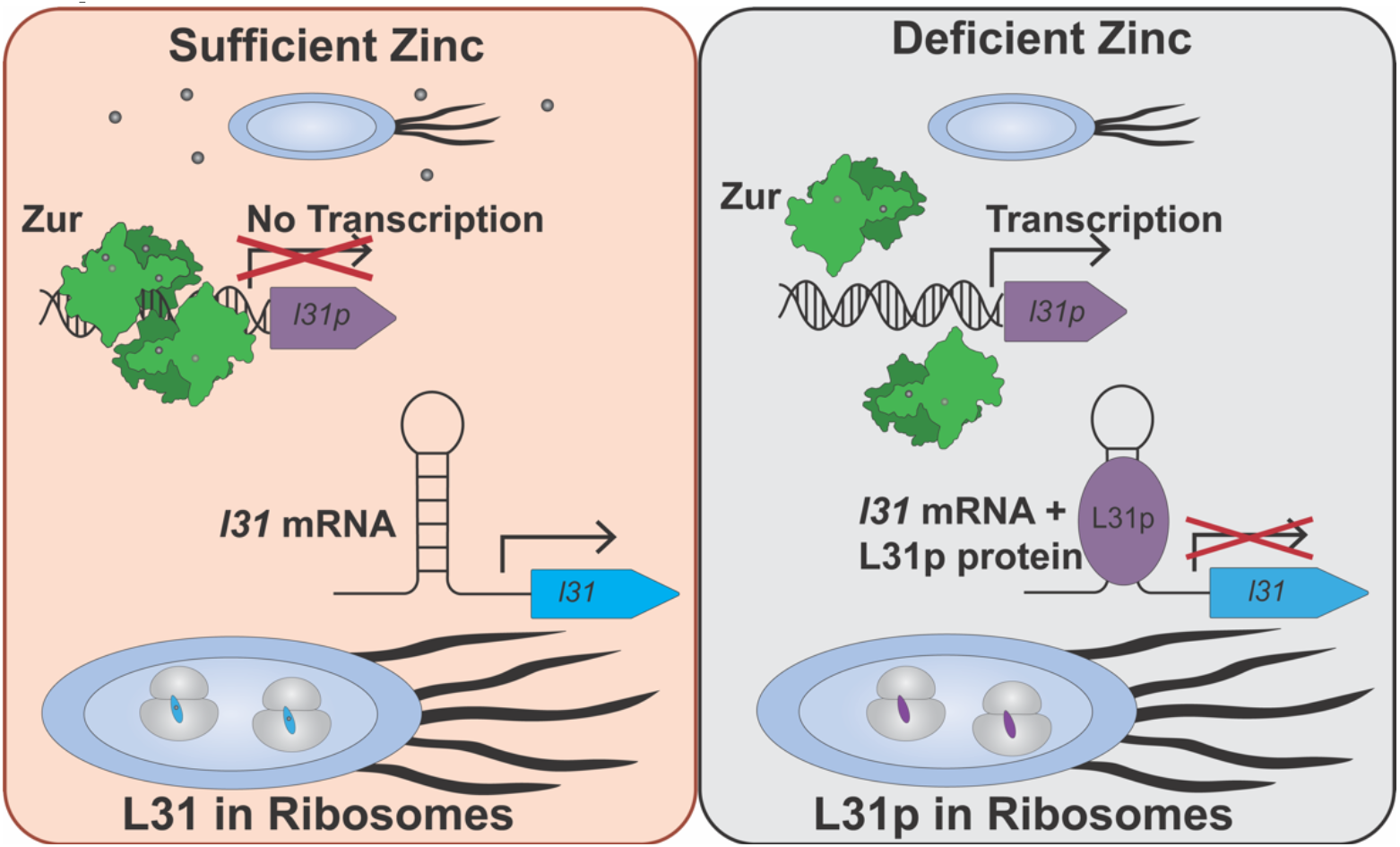

## Introduction

Zinc serves many important functions in bacteria, performing both as an enzymatic co-factor and structural roles in proteins (1, 2). Sufficient intracellular zinc levels are needed to maintain these functions, but excess intracellular zinc can lead to toxicity (3). To survive in a range of zinc concentrations in external environments and within hosts, bacteria have adapted several mechanisms to maintain intracellular zinc levels within a viable range (4, 5)

One example of zinc adaptation in bacteria is switching of ribosomal proteins from zinc-binding to zinc-lacking paralogues, which releases zinc from the zinc-binding ribosomes for other cellular functions (6, 7). The zinc-binding paralogues (called C+) have 4 conserved cysteines for binding zinc, while the zinc-lacking paralogues (called C-) lack this zinc binding motif (8). In the model organism *E. coli*, the C+ ribosomal proteins are L31 and L36 (also referred to as L31A and L36A, respectively, encoded by the genes *rpmE* and *rpmJ.*) L31 and L36 have C-paralogues, L31p and L36p (also referred to as L31B and L36B or YkgM and YkgO, encoded by the genes *ykgM* and *ykgO*)(8, 9). In the *E. coli* ribosome, L31 acts as a flexible bridge connecting the large 50S and small 30S subunit, switching between an extended and kinked conformation as the ribosome ratchets during translation elongation (10). In *l31* knockout strains, *E. coli* have decreased efficiency of 70S ribosome assembly, decreased 70S ribosome stability, decreased *in vitro* translation output, increased frameshifting, and decreased growth, especially at lower temperatures and in rich media (11–14). L36 is a small and basic ribosomal protein involved in late-stage assembly of the 50S subunit and organization of the 23S rRNA (15, 16). Recent crystallography and 2D gel electrophoresis studies of the *E. coli* ribosome indicate that L31p and L36p can replace L31 and L36 in the same general location in the ribosome, leading to ribosomes that can translate proteins with similar efficiency as with the C+ proteins present (13, 17). In addition to their dependence on intracellular zinc levels, L31p and L36p were identified in ribosomes in higher abundance during stationary phase than exponential phase (17). The zinc-lacking protein paralogues could also alter translation in a gene-depend manner, as suggested in a recent study in *Mycobacterium smegmatis* (18).

To enact the ribosomal protein switch, microbes have evolved sophisticated gene regulatory networks that are governed by zinc availability. Much of this regulation focuses on the master transcription factor Zur, which among other targets represses transcription of *l31p* and *l36p* in the *ykgMO* operon in *E. coli* (Figure 1A) (19, 20). Zur is in the Fur family of transcription factors and represses gene transcription by binding to a consensus palindromic sequence in the promoter called a Zur box (21). Besides the *ykgMO* operon, other genes repressed by Zur in *E. coli* include the ABC transporter *znuABC*, the periplasmic zinc scavenger *zinT*, and the lysozyme inhibitor *pliG* (19, 20, 22, 23). Biochemical studies indicate that *E. coli* Zur represses transcription by binding to the *znuC* promotor at free zinc concentrations in the subfemtomolar range (5). Live cell expression experiments in *Bacillus subtilis* suggest that Zur represses different genes across a range of zinc concentrations (24). While these studies all provide a clear mechanism for how Zur decreases L31p and L36p protein levels in response to zinc, they do not address how L31 or L36 could be affected by changes in zinc availability.

**Figure 1.**
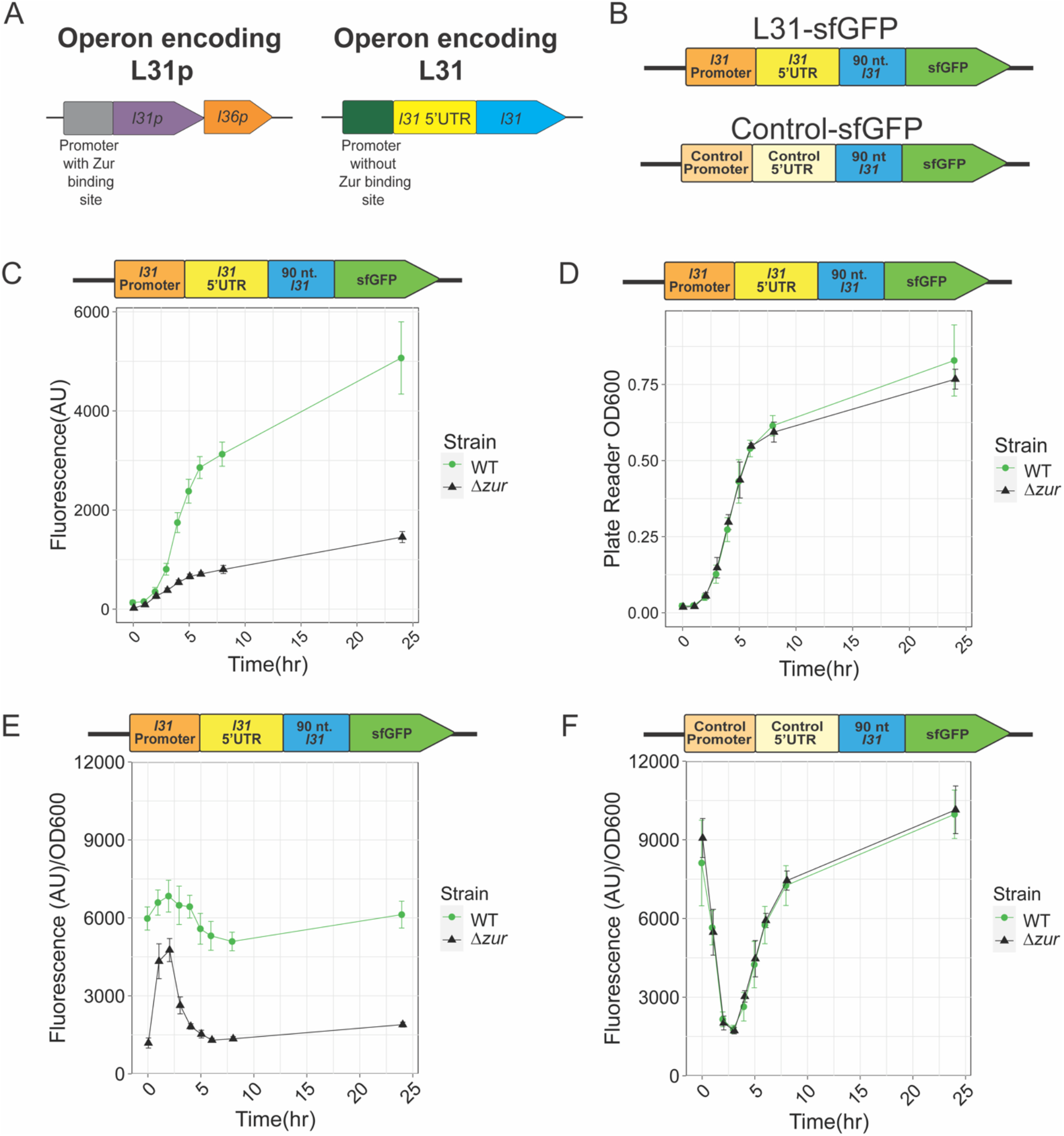
The presence of the *zur* gene increases L31-sfGFP expression in cells at all growth phases at 37 °C. A.) Operons that encode ribosomal proteins L31p and L31 in *E. coli* MG1655. B) A DNA plasmid reporter gene contains the *l31* promoter, 5’ UTR region, and a portion of the L31 coding sequence translationally fused to an sfGFP coding sequence. A control construct contains an *E. coli* sigma 70 promoter and a scrambled 5’ UTR region fused to the same coding sequence. Plasmids were transformed, grown overnight, and subcultured before measurement of fluorescence and OD_600_ in a plate reader to characterize expression from these constructs. C.) Fluorescence from L31-sfGFP plasmid in cells from 0-24 hours, measured on a plate reader D.) OD_600_ of the same samples as A., also measured directly from a plate reader. E.) sfGFP fluorescence divided by OD_600_ values from A. and B. to normalize fluorescence to cell density. F.) Normalized fluorescence/OD_600_ from control-sfGFP plasmid in cells from 0-24 hours. In each graph, the points indicate averages of three independent biological replicates, each performed with three technical replicates (cultures), for a total of nine data points (n=9). The error bars represent standard deviation of the mean.

In RNA-seq data comparing WT *E. coli* MG1655 to a *zur* knockout strain, mRNA levels of *l31* (but not *l36*) were lower when z*ur* was knocked out compared to wild-type, suggesting that Zur might regulate *l31* transcription or mRNA stability (unpublished data, Wang and O’Halloran). Unlike the previously identified Zur-regulated genes in *E. coli*, the *l31* promoter lacks the consensus Zur-binding site (20), suggesting a more complex mechanism is at play than Zur’s canonical transcriptional repression. Some bacterial species both repress and activate gene expression with Zur, but those examples still depend on Zur binding to a Zur box in the promoter. In the actinobacteria *Streptomyces coelicolor*, Zur activates expression of the zinc exporter gene *zitB* at micromolar zinc concentrations by binding at additional sites directly upstream of the main Zur box (25). In *Xanthomonas campestris*, Zur can also activate gene expression by binding to DNA in the promoter, although the inverted repeat sequence for activation differs than that for repression (26). Because the *E. coli l31* promoter region does not contain this type of sequence, we hypothesized that Zur might regulate *l31* mRNA levels expression through a different mechanism than promoter binding.

One possible explanation is that a Zur-regulated protein could alter *l31* mRNA levels through protein-DNA or protein-RNA interactions. In the recently uncovered L31 autoregulation mechanism, L31 protein is proposed to bind to the *l31* mRNA 5’ untranslated region to repress its own translation (27). Ribosomal protein autoregulation mechanisms have been identified for a series of bacterial ribosomal protein operons(28–33). A similar mechanism could explain the zinc and Zur dependent increase in *l31* mRNA in the RNA-seq data. The Zur-repressed L31p shares several structural characteristics with L31 protein, despite sharing <40% sequencing identity (17). In this model, L31p could bind the *l31* mRNA 5’UTR in a similar manner as L31. By binding to the *l31* 5’UTR, L31p protein could block L31 translation, increase mRNA decay, and/or modulate transcription. Understanding this mechanism would help explain how bacteria are able to adapt to zinc-deficient conditions, such as those presented by the host in nutritionally immunity to challenge pathogens (4, 34, 35).

### Here we address proposed mechanism

s for zinc and Zur regulation of ribosomal protein switching between L31 and its zinc-free paralogue L31p in *E. coli*. Through L31-reporter gene assays, reverse transcription quantitative polymerase chain reaction, and proteomic analysis of purified ribosomes, we find that Zur and zinc increase L31 protein and mRNA levels by repressing the repressor L31p. By connecting zinc’s regulation of L31p protein to regulation of its paralogue L31, this work proposes an RNA-protein mechanism that explains one means of bacterial adaption to zinc-deficient environments. Overall, this study increases our understanding of how bacterial ribosomes can change their composition to adapt to environmental stressors.

## Materials and Methods

### Preparation of Plasmids and Strains

Strains used in the reported assays are derivatives of *E. coli* MG1655. The strain MG1655 *Δzur* was provided by Dr. Suning Wang, and the remaining strains were created through P1 phage transduction as previously described, with BW25113 *ΔykgM* or *ΔykgO* from the Keio collection as donor strains and MG1655 wild-type or *Δzur* as recipient strains (36, 37)

Plasmids used in this manuscript are described in Table S1. Plasmids were cloned using Gibson Assembly or inverse PCR, propagated in *E. coli* TG1 competent cells in LB media, and isolated through miniprep (Qiagen.) Reporter plasmids had a p15A origin of replication, chloramphenicol resistance, and the terminator trrnB downstream of the sfGFP coding sequence. Plasmids for overexpressing ribosomal proteins *in vivo* had a ColE1 origin of replication, ampicillin resistance, the synthetic constitutive *E. coli* promoter J23108 from the Registry of Standard Biological Parts, and the terminator *trnnB* after the protein expression gene.

### Bacterial Growth Conditions

*E. coli* cells were grown in LB media (Difco LB Broth, Miller [Luria Bertani,] Fisher Scientific, Catalog #DF0446-07-5) for reporter gene and RT-qPCR assays. Antibiotic concentrations used were 34 μg/mg chloramphenicol and 100 g/mL carbenicillin (ampicillin derivative) as needed for plasmids.

Plasmid were transformed using heat shock into cells and plated on LB-agar plates with selective antibiotics. Colonies were picked and transferred to 300 μL of LB media with appropriate antibiotics, then grown at 37° C for overnight culture. Overnight cultures were then diluted 1:50 in 200 μL of fresh media, placed in 96-well culture blocks (Costar 3961 Assay block, 2 mL, 96 well standard,) covered in Breathe-EASIER covers (Diversified Biotech, cat. # BERM-2000,) and grown by shaking at 1000 RPM and 37 °C for the indicated times in a Vortemp shaker for cellular assays.

For assays that included TPEN, transformations and cultures were set up as described above. The overnight cultures were diluted in LB containing antibiotics and 100 μM TPEN (Sigma Cat. #P4413-100MG.) After 2 hours of growth, 100 μM of ZnSO_4_ (Sigma-Aldrich, cat. #221376-100G) was added to selected wells for +Zn condition. Cells were grown for an additional 2 hours with or without the additional zinc.

### Growth and Fluorescence Analysis

Unless otherwise noted, OD600 and fluorescence were measured on a Biotek Synergy H1 MF plate reader using clear-bottom 96-well plates (Thermo Scientific Nunc, #265301). Prior to plate reader measurements, 50 μL of culture was added to 50 μL of 0.1 M PBS (Sigma-Aldrich, Cat. # P4417-100TAB). Three wells with 50 μL of media in PBS were also measured to use as a blank. Fluorescence was measured with excitation or 485 nm, emission of 528 nm, gain of 50. Growth was measured simultaneously on the same plates with 600 nm absorbance.

The average fluorescence (485 nm, 528 nm) of three blanks was subtracted from the fluorescence of each well. The average OD600 of the same three blanks was subtracted from the OD600 of each sample. This calculated fluorescence value was divided by this calculated OD600 value to obtain the fluorescence/OD600 ratio to adjust for increases in fluorescence cause by increased density of cells. The average and standard deviations were calculated from the fluorescence/OD600 ratio of each sample well.

### Inductively Coupled Plasma Mass Spectrometry

400 uL of LB media was added to three metal-free tubes (VWR Centrifuge tubes, 15 mL, Ref. #525-1121,). Nitric acid (Honeywell, Fluka, 84385-2.5 L, ³0.69%, TraceSELECT for trace analysis) was added to each LB sample and then diluted in water to a final volume of 3 mL and concentration of 3% nitric acid (v/v). Samples were heated at 60 °C overnight to digest. Tubes were weighed on an analytic balance between each step. ICP-MS was performed on a computer-controlled (QTEGRA software) Thermo iCapQ ICP-MS (Thermo Fisher Scientific, Waltham, MA, USA) operating in KED mode and equipped with an ESI SC-2DX PrepFAST autosampler (Omaha, NE, USA). Internal standard was added inline using the prepFAST system and consisted of 1 ng/mL of a mixed element solution containing Bi, In, 6Li, Sc, Tb, Y (IV-ICPMS-71D from Inorganic Ventures). Each sample was acquired using 1 survey run (10 sweeps) and 3 main (peak jumping) runs (40 sweeps). The isotopes selected for analysis were 64,66Zn, and 89Y, 115In (chosen as internal standards for data interpolation and machine stability). Instrument performance is optimized daily through autotuning followed by verification via a performance report (passing manufacturer specifications). The average ppb of zinc was calculated by averaging 64Zn and 66Zn for each sample. Molarity was calculated by normalizing data to a series of standard zinc solutions (Inorganic Ventures, Christiansburg, VA, USA).”

### Ribosome Purification

The ribosome purification protocol was developed in our lab (Shepotinovskaya, Philips, *et al.*, unpublished) using a method adapted from previous literature (38). Wild type and *Δzur E. coli* MG1655 were streaked from glycerol stocks onto LB agar plates and incubated overnight at 37 °C. Cultures of one colony of WT and *Δzur* strains were grown in LB overnight at 37 °C, shaking at 250 RPM. Those cultures were used to inoculate1L of LB at 1:100 dilution for each strain. Each 1L was grown for 4-5 hours at 37 °C, with shaking at 250 rpm, to an OD_600_ of ∼0.7. The cells were harvested by centrifugation at 6000 rpm, at 4 °C, for 10 minutes. The cell pellets were resuspended in Buffer A [10 mM Tris-Cl (pH 7.4 at 4 °C), 70 mM KCl and 10 mM MgCl_2_], centrifuged at 6000 rpm, at 4 °C, for 5 minutes to remove traces of LB and then resuspended in 10 mL of fresh Buffer A. Samples were flash-frozen in liquid N_2_ and stored at −80 °C. When needed, the cells were thawed, then disrupted by sonication (60 seconds total with 1 second on/off cycles) at 30% amplitude with a Branson Digital Sonifier^®^ 450 (Branson Ultrasonics Corporation, Danbury, CT). The lysate was pre-treated with RNAse-free DNAase (20 U/mL) and CaCl_2_ (0.5 mM) and incubated on ice for 10 minutes. The lysate was then clarified from cell debris by centrifugation at 20,000 rpm, at 4 °C, for 60 minutes and used for ribosome purification.

Lysates in Buffer A were cleared through Nalgene^®^ GF-PRE and 0.45 μm SFCA syringe filters (Thermo Fisher Scientific, Waltham, MA). Ribosomes were purified with a CIMmultus™ QA-8 Advanced Composite Column [2 μm pores; 8 mL column volume (CV)] with a quaternary amine strong anion exchanger (BIA Separations d.o.o, Ajdovščina, Slovenia) using a Bio-Rad BioLogic DuoFlow™ 10 chromatography system (Bio-Rad) set to 5 mL/min flow rate. All column purification steps were performed at 4 °C. Cleared lysates were applied to the QA-8 column over 1.25 CV with 38% Buffer B (Buffer B = Buffer A + 1 M NaCl). This was followed by a wash step over 10 CV with 38% Buffer B. Ribosomes were eluted over 10 CV with 46% Buffer B. The column was cleared of any remaining protein/nucleic acids with 100% Buffer B over 5 CV. Throughout the purification, 7.5 mL fractions were collected during the injection, wash (38% Buffer B) and clearing steps (100% Buffer B), and 4.0 mL fractions were collected during the ribosome elution step (46% Buffer B). Various fractions, corresponding to chromatogram peaks (measured at 280 nm absorbance), were analyzed by SDS-PAGE for the presence of ribosomal proteins. Fractions containing ribosomes were combined, exchanged into ribosome Storage Buffer [50 mM HEPES (pH 7.5 at 4 °C), 10 mM MgCl_2_ and 70 mM NH_4_Cl], and concentrated using an Amicon Ultra-15 centrifugal filter units (3K MWCO) (MilliporeSigma, Burlington, MA) at 4 °C to a volume of 1-2 mL. Ribosome concentrations were measured using the Pierce™ BCA Protein Assay Kit (Thermo Fisher Scientific) to measure total protein concentration. The presence of L31 and L31p in the ribosome was confirmed by proteomics analysis. Proteomics services were performed by the Northwestern Proteomics Core Facility.

### TMT Sample Preparation

All reagents were purchased from Thermo Fisher Scientific unless otherwise notes. Purified ribosomes were sonicated for three rounds of 15 seconds and centrifuged at 10,000 x g for 10 minutes. Protein concentration was determined by BCA and 100 µg protein was diluted in a final volume of 200 µL with 100 mM TEAB prior to reduction with TCEP at a final concentration of 10 mM for 1 hour at 50 °C. Reduced cysteines were derivatized with iodoacetamide at a final concentration of 18.75 mM for 30 minutes in the dark. Proteins were precipitated with 8 volumes of ice-cold acetone and 1 volume TCA and incubated overnight at −20 °C. Precipitates were pelleted at 15,000 x g for 15 minutes at 4 °C, washed twice with ice-cold acetone, and air dried followed by resuspension in 100 mM TEAB. Samples were digested at 37 °C with 0.5 µg Lys-C for 6 hours then 1 µg trypsin overnight. TMT labelling and desalting were performed according to manufacturer’s recommendations (Thermo Fisher Scientific).

### LC-MS/MS Analysis for TMT

Peptides were analyzed by LC-MS/MS using a Dionex UltiMate 3000 Rapid Separation nanoLC and a Q Exactive HF Hybrid Quadrupole-Orbitrap Mass Spectrometer (Thermo Fisher Scientific Inc, San Jose, CA). Approximately 1 μg of peptide samples was loaded onto the trap column, which was 150 μm x 3 cm in-house packed with 3 μm ReproSil-Pur beads (Maisch, GmbH). The analytical column was a 75 μm x 10.5 cm PicoChip column packed with 3 μm ReproSil-Pur beads (New Objective, Inc. Woburn, MA) at 300 nL/min. Solvent A was 0.1% formic acid (FA) in water and Solvent B was 0.1% FA in acetonitrile (ACN). The peptides were separated on a 180-min analytical gradient from 5% ACN/0.1% FA to 30% ACN/0.1% FA. The mass spectrometer was operated in data-dependent mode. Data were acquired in technical duplicate. The source voltage was 2.10 kV and the capillary temperature was 320 °C. MS1 scans were acquired from 300-2000 m/z at 60,000 resolving power and automatic gain control (AGC) set to 3×10^6^. The top 15 most abundant precursor ions in each MS1 scan were selected for fragmentation. Precursors were selected with an isolation width of 2 Da with fixed first mass at 100 m/z for a reporter ion detection and fragmented by Higher-energy collisional dissociation (HCD) at 30% normalized collision energy in the HCD cell. Previously selected ions were dynamically excluded from re-selection for 20 seconds. The MS2 AGC was set to 1×10^5^.

### Proteomics Data Analysis

Proteins were identified from the tandem mass spectra extracted by Xcalibur version 4.0. MS/MS spectra were searched against the Uniprot E. Coli K12 database using Mascot search engine (Matrix Science, London, UK; version 2.5.1). All searches included carbamidomethyl cysteine as a fixed modification and oxidized Met, deamidated Asn and Gln, acetylated N-term, and TMT6-plex on Lys and N-term as variable modifications. Three missed tryptic cleavages were allowed. The MS1 precursor mass tolerance was set to 10 ppm and the MS2 tolerance was set to 0.05 Da. TMT reporter ion quantification and validation of identified peptides and proteins were performed by Scaffold software (version Scafpercent_4.8.4, Proteome Software Inc., Portland, OR). Intensity of reporter ions were calculated by Scaffold Q+. A 1% false discovery rate cutoff was applied at the peptide level. Only proteins with a minimum of two peptides above the cutoff were considered for further study. An ANOVA test with Benjamini and Hochberg false discovery rate (FDR) correction was applied to the comparison among the conditions, using a 0.05 threshold for statistical significance.

### RT-qPCR of *sfGFP* mRNA

Overnight cultures were grown in LB media in a Vortemp shaker as described above with 3 cultures per strain/plasmid combination, each from a different transformed colony. Each overnight culture was diluted 1:50 in LB-antibiotic the next morning into 3 separate 200 μL cultures to increase the volume for analysis while still allowing for sufficient oxygenation of each well. Cells were then grown as described in “Bacterial Growth Conditions” above. After 2 hours, or early exponential phase, 150 μL of each culture from samples of the same strain/plasmid combination were combined and pelleted by centrifugation. RNA was isolated by resuspending in Trizol (Ambion, Ref #15596018) extracting with chloroform (Acros Organics, Code 423555000) and ethanol precipitation. RNA was quantified using a Qubit ssRNA broad range assay (Catalog # Q10211). DNA from the samples was digested by incubating 200 ng of total RNA from each sample with DNAse Turbo (Invitrogen, Ref #AM2238) in Turbo DNAse buffer (Invitrogen, #4022G) for 1 hour at 37 °C to digest. Phenol:chloroform extraction was then performed (Acros Organics Phenol/Chloroform/Isoamyl alcohol (25:24:1) Code 327115000; Acros Organics Chloroform, Code 423555000) followed by an additional ethanol precipitation. RNA was quantified again using a Qubit ssRNA high sensitivity assay to quantify the volume needed for the reverse transcription (RT) reaction (Catalog #Q32852). Reverse transcription was then performed by incubating 1 ng of RNA with 0.5 μL of 10 mM dNTPs and 0.5 μL of 2 μM reverse transcription primer (Table S3) in a 6.5 μL volume at 65 °C for 5 minutes, then placed on ice for 5 minutes. For each sample, one reaction was performed with the sfGFP reverse transcription primer, and the other was performed with the 16S rRNA primer as a control. To those reactions, 2 μL of First Strand buffer (Invitrogen, P/N Y02321,) 0.5 μL of 0.1 M fresh DTT, 0.5 μL of RNAse-out (Thermo Fisher Scientific, cat. # 10777019,) and 0.25 μL Superscript III reverse transcriptase (Invitrogen, P/N 56575) were added to a volume of 10 μL. Samples were then incubated at 55 °C for 1 hour. Control reactions without Superscript III with each reverse transcription primer were performed to verify the absence of plasmid DNA contamination.

Quantitative PCR was performed on the above cDNA samples in technical triplicate, performing the PCR reaction in three separate wells for the same reverse transcription sample. Each reaction consisted of 1 μL of reverse transcription samples, 5 μL SYBR Green mastermix (Applied Biosystems by Thermo Fisher Scientific, Ref. #4344463) and 0.5 μL of 2 μM of each corresponding forward and reverse qPCR primers: sfGFP qPCR primers for sfGFP RT reactions, and 16S rRNA qPCR primers for the 16S rRNA RT reactions (Table S3). Control reactions without reverse transcription enzyme from above, and reactions with nuclease-free water instead of cDNA template were run simultaneously. Samples were prepared on a 96-well PCR plates (Corning, Thermowell gold PCR plates, polypropylene, Ref. #3752) and covered with a clear sealing cover (Thermo Scientific, Ref. #23701). PCR was performed on a BioRad CFX Connect Real-Time System with the following cycling settings: 50 °C for 2 minutes, 95 °C for 10 minutes, 40 cycles of 95°C for 15 seconds and 60 °C for 1 minute, followed by a melt curve analysis from 65 to 95 °C in 0.5 °C increments. The cycle threshold, or C_q_, for each sample was calculated by the software CFX Manager Version 3.1.1517.0823. The Δ C_q_ for each sample was calculated by subtracting 16S rRNA average C_q_ from each sfGFP C_q_ value from the same RNA isolation sample (i.e. the C_q_ with sfGFP primers minus the C_q_ with 16S rRNA primers, both from WT strains with the control-sfGFP plasmid.) The average Δ C_q_ of the WT control-sfGFP sample was subtracted from the Δ C_q_ of each well to obtain the ΔΔC_q_ of each well. Relative quantification of each well was calculated as 2^(- ΔΔC_q_.)

### Prediction and Design of 5’UTR Structures

RNA structures were predicted using the online software NUPACK (39–43). For the *l31* 5’UTR and its mutants, RNA from the transcription start site to the 10^th^ nucleotide of the *l31* coding region were including in the input.

The scrambled control was generated by adding a ribosome binding site and randomly scrambling the remaining nucleotides to generate a new 5’UTR that has the same length and nucleotide composition as the *l31* 5’UTR. Mutations to the 5’UTR were designed by trial-and-error and input in NUPACK to design mutants in which the minimum free energy structure only changes in the targeted location or does not change, depending on the experiment.

### Modeling Protein-RNA Interactions

Images were rendered on the software ChimeraX using PDB 6i7v (17, 44, 45).

### Material Availability

All plasmids are deposited at Addgene depository, Deposit # 80991. Plasmids are described in Table S1.

## Results

### The presence of zur increases L31-reporter levels

Zur is well known as a zinc-responsive repressor of transcription in bacteria (20, 21, 46) In a previous un-published RNA-Seq study comparing transcriptome wide gene expression in WT and *Δzur E. coli* cells, we found evidence that Zur could potentially up-regulate *l31* mRNA levels (Wang and O’Halloran, unpublished data.) We first sought to corroborate this finding using reporter gene assays. To determine how the presence of Zur, combined with zinc, could affect L31 protein levels, we constructed a sfGFP reporter plasmid by fusing 400 nucleotides of the *l31* promoter region, the 105 nt-long *l31* 5’untranslated region (5’UTR), and the first 90 nt of the *l31* coding region (encoding 30 of L31’s 70 amino acids) translationally fused to the superfolder sfGFP coding sequence (Figure 1B). This expression construct was then placed on a p15A plasmid with chloramphenicol resistance (47). Similarly, we constructed a control DNA construct by fusing an *E. coli* sigma70 constitutive promoter, a scrambled *l31* 5’UTR, and the same fusion protein coding sequence on the same plasmid backbone (Figure 1B).

We next performed gene expression assays by transforming these plasmids in both WT and *Δzur* MG1655 cells, growing colonies in LB media overnight, subculturing, and characterizing sfGFP fluorescence and cell growth over time with a plate reader (Figure 1C-F, S1A). The fluorescence from sfGFP for each culture was divided by its plate reader-measured absorbance at 600 nm (OD_600_) in order to normalize for cell density (Figure 1E). The OD_600_ values were measured by diluting cultures 2x and measuring on a plate reader, which gave similar trends as when measured using a standard cuvette (Figure 1D, S1B, S1C). At all measured timepoints, expression of the L31-sfGFP construct was greater in WT than *Δzur* cells (Figure 1E). In addition, the control plasmid showed similar levers of fluorescence/OD_600_ in both WT and *Δzur* cells (Figure 1F, S1D-F). Taken together, these results show that the presence of *zur* results in increased expression from the *l31* expression context and suggest a role for Zur in up-regulating L31 expression.

### Exogenous zinc further increases L31-reporter levels in the presence of zur

Because Zur is a zinc-responsive transcriptional repressor of a variety of *E. coli* genes, we next sought to determine if L31-sfGFP protein levels were also affected by exogenous zinc concentration in the media. The average zinc concentration in the batch of LB media used in these experiments was established by ICP-MS to be 91.5 μM (Table S2). To compare zinc-limitation and zinc replete conditions, 100 μM TPEN was added to LB media in the subcultures. To half of these samples, 100 μM zinc was later added, and growth was analyzed as described above (see Materials and Methods). In WT cells transformed with the L31-sfGFP construct, we observed a significant increase in fluorescence/OD_600_ levels in the zinc-replete relative to zinc-limitation conditions (Figure 2A). However, the same construct transformed in the *Δzur* strain showed no significant dependence on zinc availability (Figure 2A). Parallel experiments in strains containing the control plasmid showed a small *decrease* in fluorescence/OD_600_ in zinc-replete conditions regardless of the presence of *zur* (Figure 2B). Overall, these results show that zinc leads to increased expression of the regulated *l31-sfGFP* expression construct, but only in the presence of *zur*. This suggests that that both Zur and zinc together increase L31 expression.

**Figure 2.**
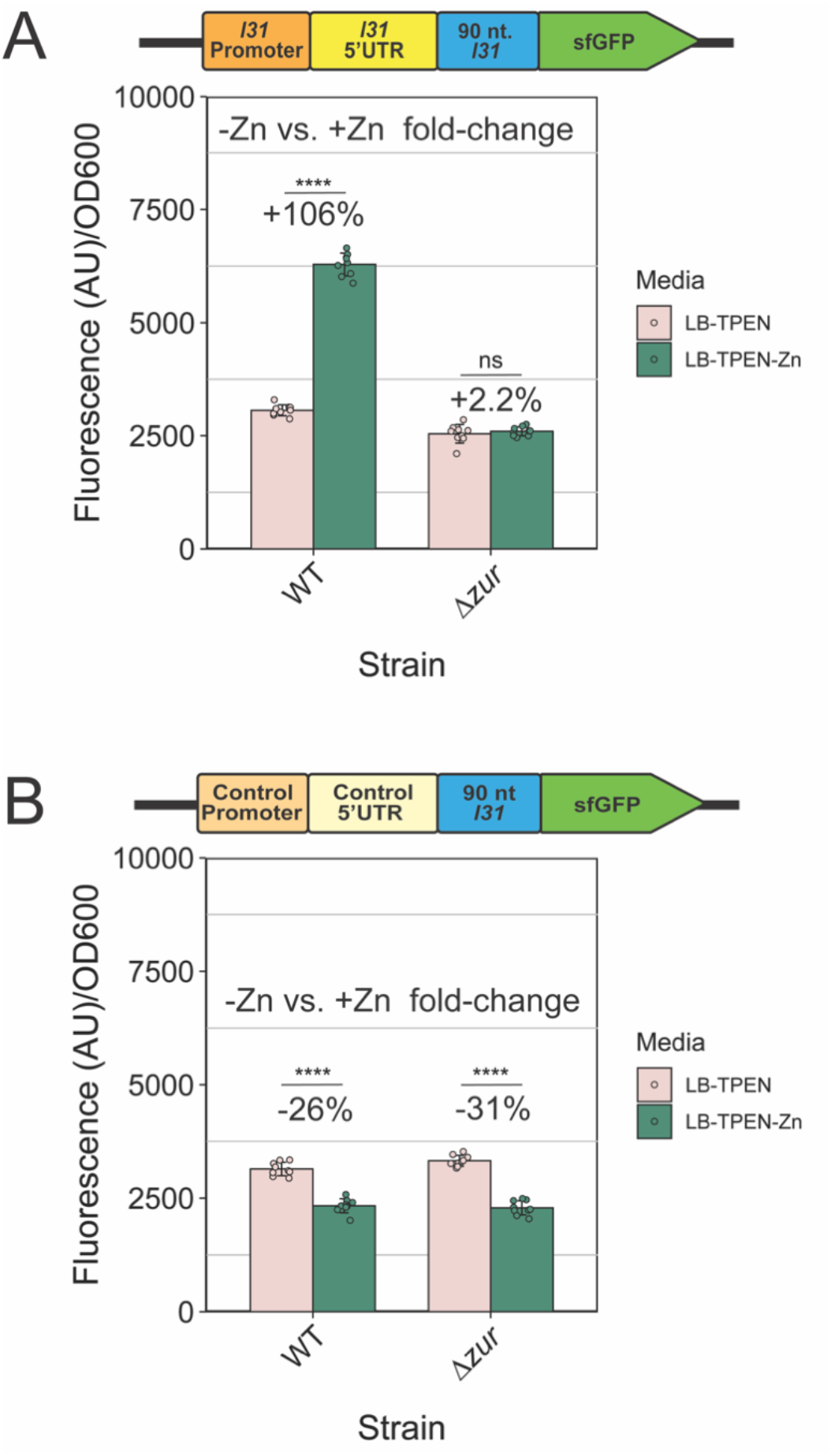
Zinc increases L31-sfGFP expression from regulated constructs in cells when *zur* is present. A.) Fluorescence divided by OD_600_ of subculture cells with the L31-sfGFP reporter plasmid grown in LB + 100 μM of the zinc chelator TPEN for 2 hours, plus an additional 2 hours with or without the addition of 100 μM ZnSO_4_ before plate reader data collection B.) Fluorescence divided by OD_600_ of subculture cells with the control-sfGFP reporter plasmid measured in the same way. In each graph, the bars indicate averages of three independent biological replicates, each performed with three technical replicates (cultures) for a total of nine data points (n=9). The error bars represent standard deviation of the mean. Percent change was calculated using the average fluorescence/OD_600_ in the equation 100*(*+*Zn – No Zn) / No Zn. Significance was calculated with a 2-tailed student’s t-test between the fluorescence/OD_600_ values for each group. p-value < 0.05 = *, p-value < 0.01 = **, p-value < 0.001 = ***, p-value < 0.0001 = ****.

### zur increases l31 mRNA levels and L31 incorporation into ribosomes

To determine if the presence of *zur* also increases the expression of *l31-sfGFP* at the mRNA level, we performed qRT-PCR on *E. coli* WT and *Δzur* subcultures. These subcultures were grown for two hours to early exponential phase using the same strains, plasmids, and growth methods used in the L31-sfGFP *in vivo* expression assays. Similar to our previous sfGFP fluorescence results, the *l31-sfGFP* mRNA was shown to be higher in the presence of *zur*, but only for the plasmid with the *l31* promoter and 5’UTR (Figure 3A, Figure S2.)

**Figure 3.**
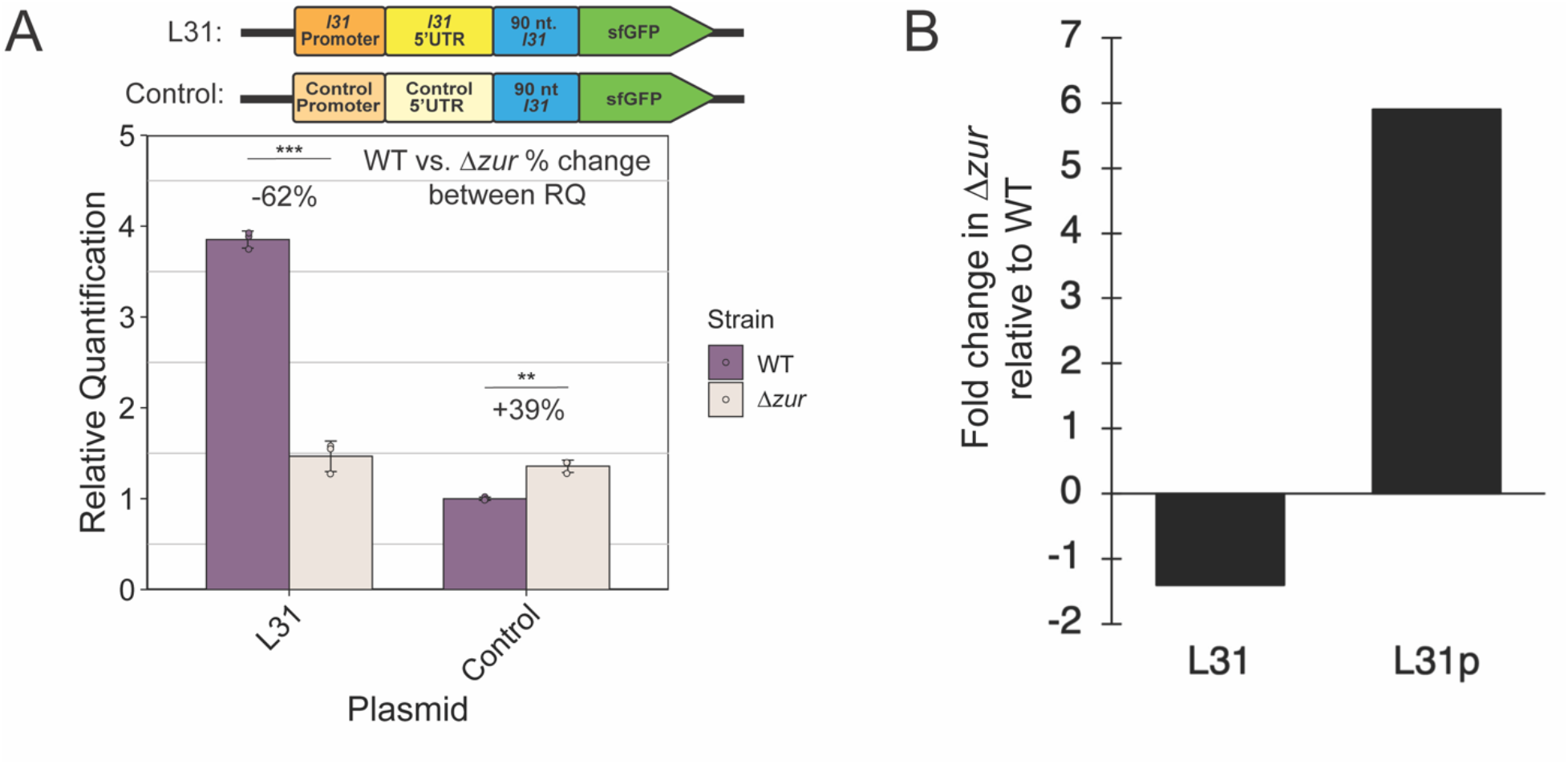
The presence of *zur* increases the levels of *l31-sfGFP* mRNA and L31 incorporation into the ribosome. A.) Reverse-transcription quantitative PCR of *l31-sfGFP* mRNA in WT and *Δzur E. coli* with L31-sfGFP and control-sfGFP plasmids in LB. The bars indicate averages of three technical replicates (independent wells of the same qPCR reaction) of one experiment. See Materials and Methods for experimental details. A second biological replicate of this experiment is shown in Figure S2. The error bars represent standard deviation of the mean. Percent change was calculated using the average fluorescence/OD600 values in the equation 100*(*Δzur* – WT) / WT. B.) Ribosomes were purified from cells grown in LB to exponential phase OD_600_ = 0.7. L31 and L31p protein content was measured using mass spectrometry on TMT-labelled ribosomes. Fold-change of L31 and L31p protein quantity in *Δzur* ribosomes was calculated compared to WT ribosomes.

We next investigated how the presence of *zur* affects the abundance of L31 vs. L31p protein that are incorporated into cellular ribosomes. A recent study in *Mycobacteria smegmatis* suggests that alternative zinc-lacking ribosomal protein paralogues could alter ribosome translation gene-specifically, indicating a potential role of ribosomal protein paralogue switching in modulating translation to adapt to environmental stressors (18). The first step to study a possible similar mechanism in *E. coli* is to determine if the ribosomal protein L31p replaces L31 when Zur is de-repressed (or *zur* is absent). To investigate this, ribosomes were purified from both WT and *Δzur* strains grown in exponential phase cultures in LB media from diluted overnight cultures (Figure S3), avoiding a possible increase in L31p resulting from stationary phase (17). Mass spectrometry was performed on both ribosome samples to determine the change in L31 and L31p from WT to *Δzur* ribosomes. Based on this analysis, we found that ribosomal incorporation of L31 was lower and L31p incorporation higher in *Δzur* cells, suggesting that a decrease in L31 protein levels in *Δzur* cells also results in decreased L31 in the ribosome (Figure 3B). These results demonstrate that the absence of *zur* can lead to changes in *l31* mRNA levels, which could lead to increases in L31p and decreases of L31 incorporation into cellular ribosomes.

### L31p is required for zur and zinc to increase L31-reporter levels

We then aimed to determine the general mechanism by which Zur and zinc act to increase L31 protein expression. Zinc-loaded Zur is known to be a repressor of transcription of the *ykgMO* operon by binding to the Zur box in the promoter of the operon (19, 20). Recent work suggests that the L31 protein represses its own translation by binding to its mRNA (27). We therefore hypothesized that L31p, which is structurally similar to L31 and encoded in the *ykgMO* operon, could act as a repressor of L31 translation when that operon is de-repressed by Zur. To determine if Zur and zinc act to increase L31 protein levels through the Zur-regulated ribosomal protein L31p (or L36p, also encoded in the *ykgMO* operon), we first generated strains lacking these genes, with or without *zur*. We did not knock out *l31* to avoid growth defects of Δ*l31* (Δ*rpmE*) (12), which could confound results with inconsistencies in growth phase. *These* strains were transformed with the L31-sfGFP and control-sfGFP plasmids, grown in LB cultures overnight, subcultured for various timepoints, and characterized for fluorescence and OD_600_ on a plate reader (Figure S4A-D). Comparison of fluorescence/OD_600_ between *zur-*containing and *Δzur* strains of the three different contexts (WT, *Δl36p*, *Δl3lp*) at 6 hours of subculturing allowed us to investigate how *l36p* and *l31p* affect *zur*-mediated regulation. For the regulated L31-sfGFP construct in both the WT and *Δl36p* contexts, we observed a decrease in expression with *Δzur* as before (Figure 4A, S4E-F), but not with the control plasmid (Figure 4B, S4G-H), suggesting that *l36p* does not play a principal role in regulation of L31-sfGFP. In contrast, the *Δl31p* context showed no repression of the L31-sfGFP construct when knocking out *zur* (Figure 4A). These data suggest that knocking our *zur* only decreases L31-sfGFP levels in the presence of *l31p*.

**Figure 4.**
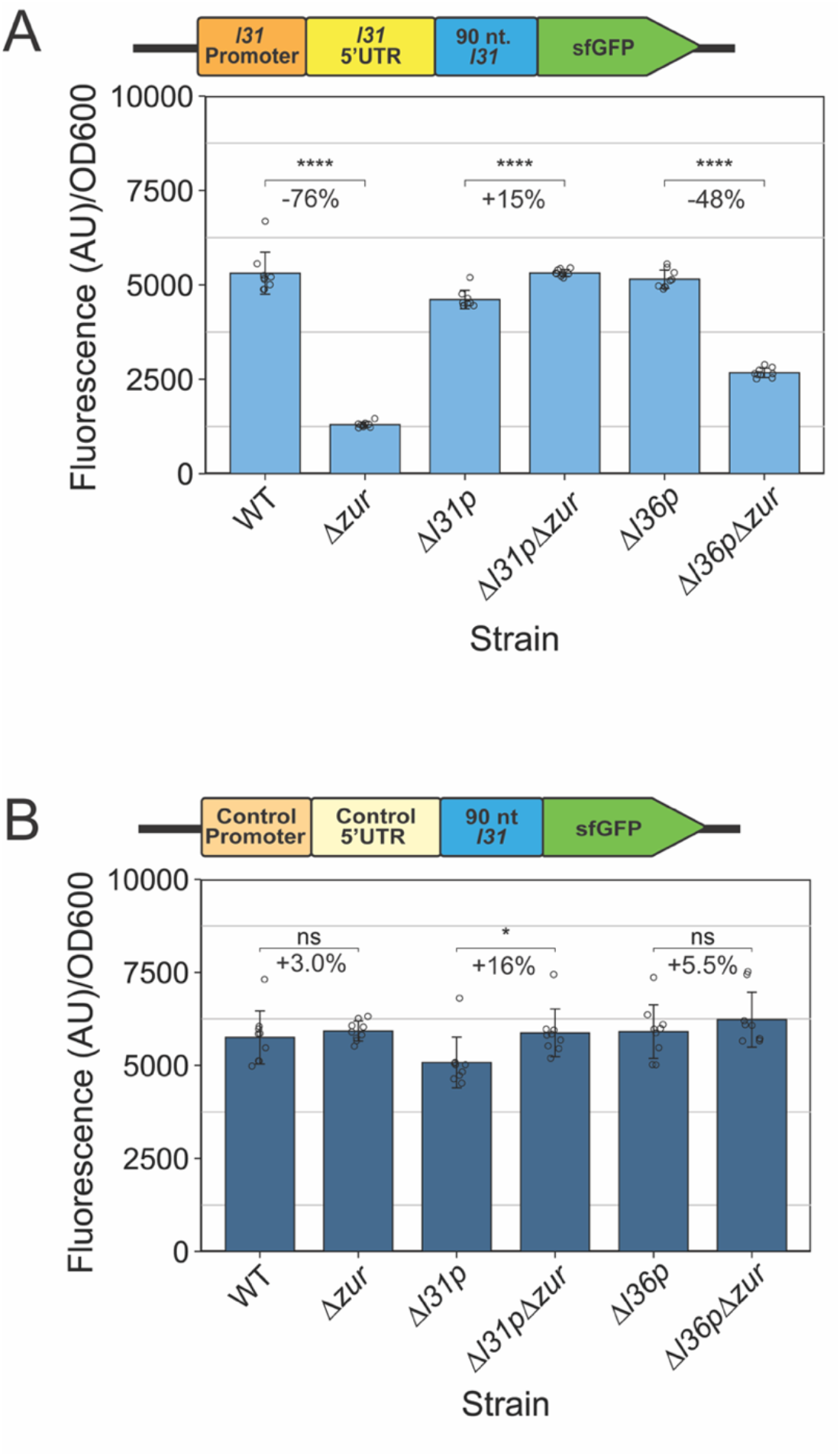
Regulation of L31-sfGFP expression by *zur* requires L31p. A.) L31-sfGFP fluorescence divided by OD_600_ measured on a plate reader for different strains grown in LB for 6 hours. B.) Control-sfGFP fluorescence divided by OD_600_ on plate reader for different strains grown in LB for 6 hours. In each graph, the bars indicate averages of three biological replicates (independent experiments), each performed with three technical replicates (cultures per experiment) for a total of nine data points (n=9). The error bars represent standard deviation of the mean. Percent change between pairs of bars was calculated using the average fluorescence/OD_600_ values in the equations 100*(*Δzur* – WT) / WT, 100*(*Δl31pΔzur* – *Δl31p*) / *Δl31p*, and 100*(*Δl36p Δzur* – *Δl36p*) / *Δl36p*, respectively. Significance was calculated with a 2-tailed student’s t-test between the fluorescence/OD_600_ values for each comparison group. p-value < 0.05 = *, p-value < 0.01 = **, p-value < 0.001 = ***, p-value < 0.0001 = ****.

To determine if L31p’s repression is also regulated by zinc, the same strains were tested for sfGFP fluorescence in the presence of 100 μM TPEN, with or without added 100 μM zinc (Figure 5, S5). Comparing the observed fluorescence/OD_600_ measurements in the zinc and no zinc conditions between strains showed that the *Δl31p* context broke Zn-mediated regulation, while the *Δl36p* context still showed a Zn-mediated increase in expression though with quantitatively different levels (Figure 5A). This trend was not observed with the control-sfGFP plasmid (Figure 5B).

Combined, these data further support that Zur and L31p - but likely not L36p - are elements in the Zur regulatory circuitry that increases L31 expression in sufficient zinc. Because Zur and zinc were previously reported to repress *l31p* expression (19), we hypothesize that L31p could be a Zur-regulated repressor of L31 protein expression. By playing the role as a translational repressor, the role of L31p is to flip the repression logic of Zur into an activator.

**Figure 5.**
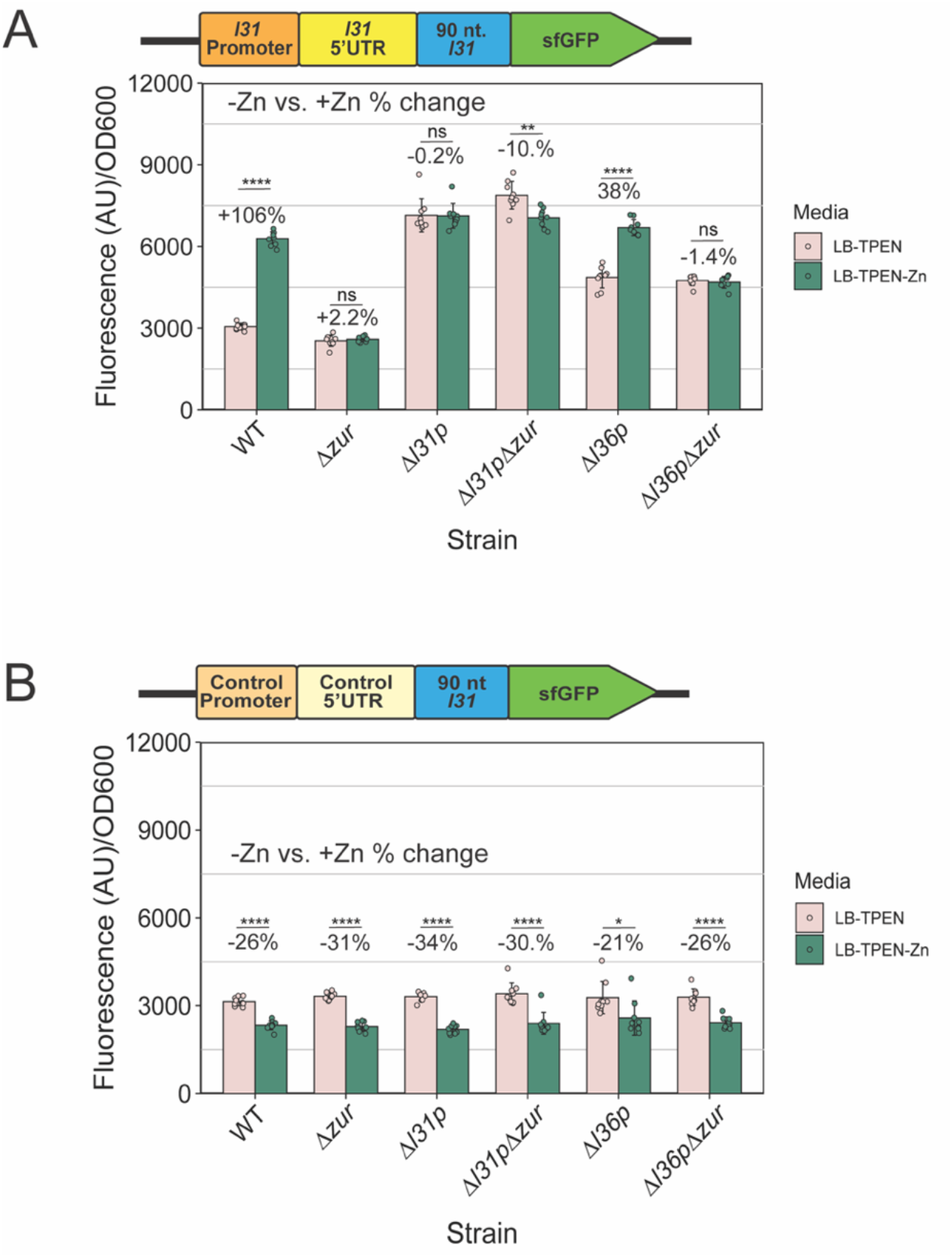
L31p is necessary for zinc to increase L31-sfGFP expression in cells. A.) Fluorescence/OD_600_ with L31-sfGFP plasmid in various knockout strains, grown in LB + 100 μM of TPEN for 2 hours, then with or without the addition of 100 μM ZnSO_4_ for an additional 2 hours. B.) Fluorescence/OD_600_ with control-sfGFP plasmid in various strains, grown in LB + 100 μM of TPEN for 2 hours, then with or without the addition of 100 μM ZnSO_4_ for an additional 2 hours. Note: the results on the WT and *Δzur* strains shown in this figure are also shown in Figure 2A-B. In each graph, the points indicate averages of three biological replicates (independent experiments), each performed with three technical replicates (cultures per experiment) for a total of nine data points (n=9.) The error bars represent standard deviation of the mean. Percent change was calculated using the average fluorescence/OD_600_ values in the equation 100*(*+*Zn – No Zn) / No Zn for each strain. Significance was calculated with a 2-tailed student’s t-test between the fluorescence/OD_600_ values for no added zinc vs added zinc for each strain. p-value < 0.05 = *, p-value < 0.01 = **, p-value < 0.001 = ***, p-value < 0.0001 = ****. *A stem-loop in the l31 mRNA 5’UTR is required for the* zur*-dependent increase in L31-reporter output*

The L31 protein can autoregulate its own translation through a mechanism that requires the 5’UTR of its mRNA for repression (27). We therefore hypothesized that the 5’UTR region of the *l31* mRNA is similarly required for *zur* to increase L31-sfGFP levels in our assays. To test this hypothesis, we constructed reporter plasmids that contained either the *l31* 5’UTR or a scrambled 5’UTR plus a ribosome binding site, in combination with either the *l31* promoter region or a synthetic sigma 70 promoter (Figure 6A). When WT and *Δzur* cells were grown in LB with these plasmids, the *Δzur* cells showed a significant decrease in sfGFP expression only when the constructs contained the *l31* 5’UTR, independent of promoter region (Figure 6B). This result demonstrated that the *l31* 5’UTR—not the promoter—is required for *zur*’s regulation of L31 protein levels.

**Figure 6.**
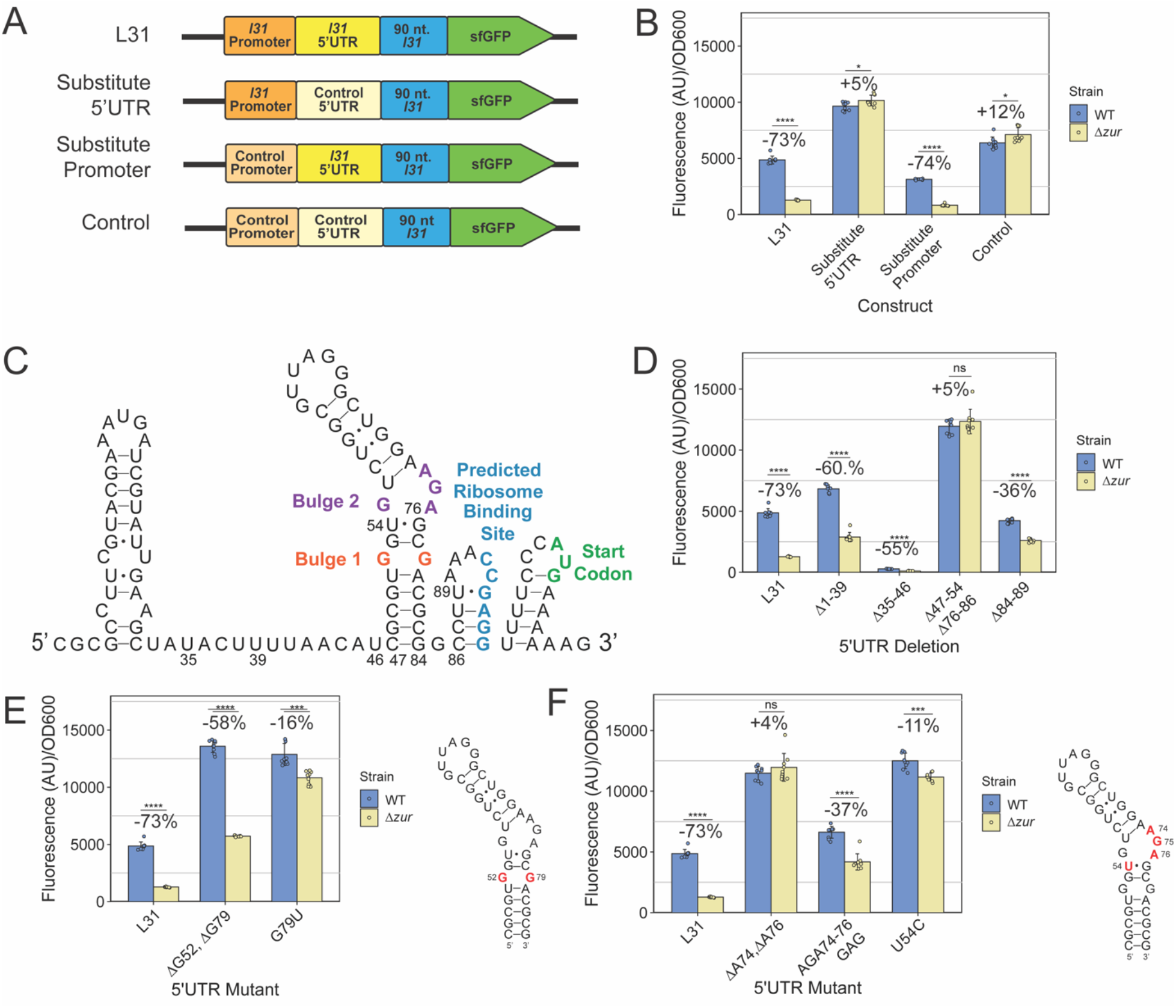
The asymmetric inner loop in the *l31* 5’UTR stem loop 2 is important for *zur*’s regulation of L31-sfGFP expression. A.) DNA constructs on plasmids used in promoter and 5’UTR substitution experiments. B.) Measured Fluorescence/OD_600_ with promoter and 5’UTR substitutions on reporter plasmids. C.) Predicted structure of the *l31* 5’UTR mRNA, drawn using secondary structure obtained from the software NUPACK. The predicted ribosome binding site and start codon are annotated. D.) Fluorescence/OD_600_ with L31-sfGFP plasmids with 5’UTR short region deletions E.) Fluorescence/OD_600_ with L31-sfGFP plasmid with mutations to the *l31* 5’UTR Bulge 1 (G52 & G79,) and F.) Bulge 2 (AGA74-76.) In each graph, the bars indicate averages of three biological replicates (independent experiments), each performed with three technical replicates (cultures per experiment) for a total of nine data points (n=9.) The error bars represent standard deviation of the mean. Percent change was calculated using the average fluorescence/OD600 values in the equation 100*(*Δzur* – WT) / WT for each construct. Significance was calculated with a 2-tailed student’s t-test between the fluorescence/OD_600_ values for WT and *Δzur* with the same plasmid. p-value < 0.05 = *, p-value < 0.01 = **, p-value < 0.001 = ***, p-value < 0.0001 = ****.

We next asked which sequence or structural features of the *l31* 5’UTR may be responsible for this regulation. The *l31* 5’UTR sequence is highly conserved and is predicted to fold into a secondary structure consisting of 4 stem-loops, including a long, asymmetrical stem-loop (Figure 6C) (27, 48) Using this secondary structural model as a guide and NUPACK RNA structure prediction software, we mutated regions of the *l31* 5’UTR within our expression plasmid constructs and performed gene expression assays to determine which structural features are most important in this regulatory mechanism (Figure S6) (39–43). Deletion of the first 39 nucleotides showed similar L31-sfGFP expression trends to WT (Figure 6D). Deleting part of the predicted single stranded region between the first and second stem-loop (*Δ*35-46) caused low overall expression — a result seen in L31 autoregulation — but still a decrease from *Δzur* vs. WT (Figure 6D) (27). Deleting part of the longest stem-loop (Δ47-54, Δ76-86), however, eliminated *zur*’s ability to regulate this mechanism (Figure 6D). Deleting part of the stem-loop that binds with the RBS (Δ84-89) still allows *zur* to regulate L31-sfGFP to some extent, suggesting that the mechanism likely does not require interactions that directly block or free the ribosome binding site (Figure 6D).

We then performed a more refined mutational analysis of the long stem-loop to determine which nucleotides or sections of the stem-loop are required for regulation. When the lower predicted inner loop was deleted (ΔG52, ΔG79), *Δzur* reduced L31-sfGFP with almost as great of percent change as WT, albeit higher overall sfGFP levels (Figure 6E). However, when the predicted inner loop was closed by creating a wobble base pair (G79U), the ability of *Δzur* to repress was almost completely eliminated (Figure 6E). In the upper inner loop, deleting the 2 nucleotides A74 and A76 to symmetrize the inner loop structure fully eliminated *zur*’s regulation and resulted in higher overall expression (Figure 6F). Similar results were observed in the U54C mutation, which strengthens a wobble pair to form a stronger Watson-Crick-Franklin base pair in this region. In contrast, mutating nucleotides 74-76 while maintaining the same minimum free energy predicted structure (AGA 74-76 GAG), only reduced the percent change of *Δzur*’s repression approximately in half (Figure 6F, S6). Similarly, mutating the wobble pairs to Watson-Crick-Franklin pairs in the upper stem (U58C, U70C) had no discernable effect of regulation, nor did shrinking the upper loop (ΔG62, ΔG67) have a large effect on the percent change between strains, other than higher overall expression (Figure S7).

Overall, these data demonstrate the importance of the RNA hairpin structure within *l31* 5’UTR and, in particular, of the asymmetrical inner loop region in the ability of *zur* to upregulate L31 protein expression.

### The L31p N-terminus is needed to repress L31-sfGFP expression

Next, we examined the effect of overexpressing L31p variants in cells to determine which region of L31p is most important for L31-sfGFP regulation. To do so, we performed assays in which WT *E. coli* were grown as in previous experiments with the addition of plasmids that constitutively express L31p, mutants of L31p, or other proteins as controls. The previous study on L31 autorepression determined that the deletion of the N-terminus, but not deletion of the C-terminus, reduced L31 autorepression (27). An existing crystal structure of L31p protein within the *E. coli* ribosome showed that 3 residues of the L31p N-terminus interacts with the 5s rRNA, suggesting a possible role in the L31p N-terminus in binding to *l31* mRNA (Figure 7A-C).

**Figure 7.**
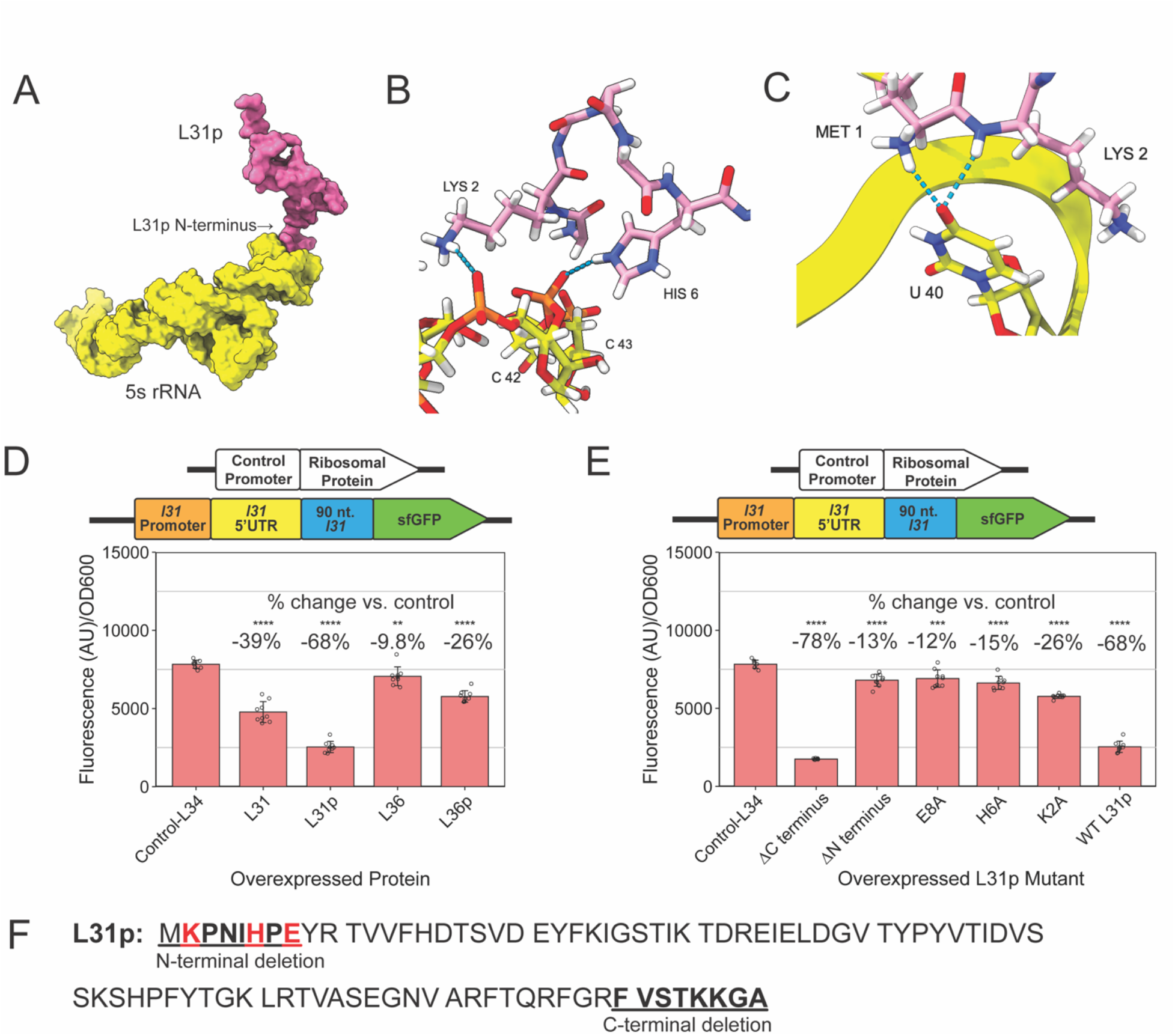
The L31p N-terminus is needed to repress L31-sfGFP expression. A.) Interaction between L31p (pink) and the 5S rRNA (yellow) in the *E. coli* ribosome. Image rendered on ChimeraX from PDB 6i7v. B. & C) Close up on polar bonds of the structure shown in A. The three L31p residues--Met1, Lys2, and His6—that interact with the 5S rRNA are shown in stick view and colored pink. D.) Fluorescence/OD_600_ with L31-sfGFP plasmid in WT *E. coli* with additional plasmid expressing different ribosomal proteins and E) L31p mutants. F.) L31p protein sequence from Ecocyc database. N- and C-terminal deleted residues are underlined, and mutated residues are shown in red. In each graph, the bars indicate averages of three biological replicates (independent experiments), each performed with three technical replicates (cultures per experiment) for a total of nine data points (n=9.) The error bars represent standard deviation of the mean. Percent change in D. and E. was calculated using the average fluorescence/OD600 values in the equation 100*(Result – Control(L34)) / Control(L34). Significance was calculated with a 2-tailed student’s t-test between the fluorescence/OD_600_ values compared to the sample expressing the control-L34. p-value < 0.05 = *, p-value < 0.01 = **, p-value < 0.001 = ***, p-value < 0.0001 = ****.

We first determined the effect of overexpressing L31p *in vivo* on L31-sfGFP levels. Compared to expression of the control protein L34, L31p decreased L31-sfGFP fluorescence with a 67% decrease, supporting its role as a repressor of L31 (Figure 7D S8A, S8B). Expression of L31, a known autorepressor, decreased its own expression with a 39% percent decrease (Figure 7D). Expressing L31p and L31 from plasmids did not decrease sfGFP expression in the control construct (Figure S8C-E). Expression of L36p also resulted in some repression of L31-sfGFP--a 24% decrease, but it is unclear from this result if L36p can also contribute to L31p’s repression of L31 protein expression (Figure 7D).

We then studied the role of different regions of L31p protein on its ability to repress L31-sfGFP. Deletion of N-terminal amino acids 2-8 prevented L31p from repressing L31-sfGFP, while deletion of C-terminal amino acids allowed L31p to repress L31-sfGFP just as effectively (Figure 7E, S8F). To evaluate roles of individual amino acids in the repression mechanisms, we focused on individual charged residues (K2, H6, E8) that have the potential to affect L31p interactions with the negatively charged *l31* 5’UTR mRNA. Independently mutating K2, H6, or E8 to an alanine each resulted in as much or almost as great of L31-sfGFP expression as deleting the N-terminus (Figure 7E, 7F, S8F). This suggests that either all these charged residues are important for the protein-RNA interaction, or these individual mutations cause too great of a structural change for L31p to bind to the *l31* mRNA.

Collectively, these results demonstrate that L31p, like L31, requires its N-terminus to repress L31 protein expression.

## Discussion

Taken together, the data in this study lead us to propose a mechanism to explain the switch to the zinc-lacking ribosomal protein L31p in zinc-deficient conditions from the zinc-binding L31 in zinc-sufficient conditions (Figure 8). In zinc-deficient conditions, the transcription factor Zur does not repress the *ykgMO* operon, allowing L31p protein to be expressed. We propose that the L31p protein in turn interacts with an RNA hairpin structure in the *l31* 5’UTR mRNA, blocking L31 translation, decreasing transcription, and/or increasing *l31* mRNA decay. In zinc-sufficient conditions, Zur represses the expression of *l31p*, preventing L31p protein from repressing L31. Results from *in vivo* L31-sfGFP assays support this mechanism by demonstrating that *l31p* is needed for *zur* and zinc to regulate L31-sfGFP expression (Figures 4A, 5A). This mechanism is further supported by increased repression of L31-sfGFP when L31p is constitutively expressed off a plasmid (Figure 7D). We identified three levels at which Zur regulates *l31*: *l31* mRNA through RT-qPCR, L31-sfGFP protein through *in vivo* reporter gene assays, and L31 protein incorporation into the ribosome through proteomic analysis of purified ribosomes (Figure 1E, 3A, 3B). The proteomics results suggest that Zur—and likely its ligand zinc— not only affect L31 protein levels within the cell but also alters the content of the ribosome, corroborating a recent examination of an *E. coli Δzur* ribosome though 2D gel electrophoresis (13). In summary, zinc-loaded Zur increases L31 protein expression by repressing a repressor of L31 expression: L31p. Similar repression of a repressor mechanisms have previously been reported with bacterial sRNAs (49). One such example is the activation of genes by the iron-sensing transcription factor Fur; Fur represses expression the small RNA RyhB, which in turn represses numerous genes by binding to their mRNA (50, 51)

**Figure 8.**
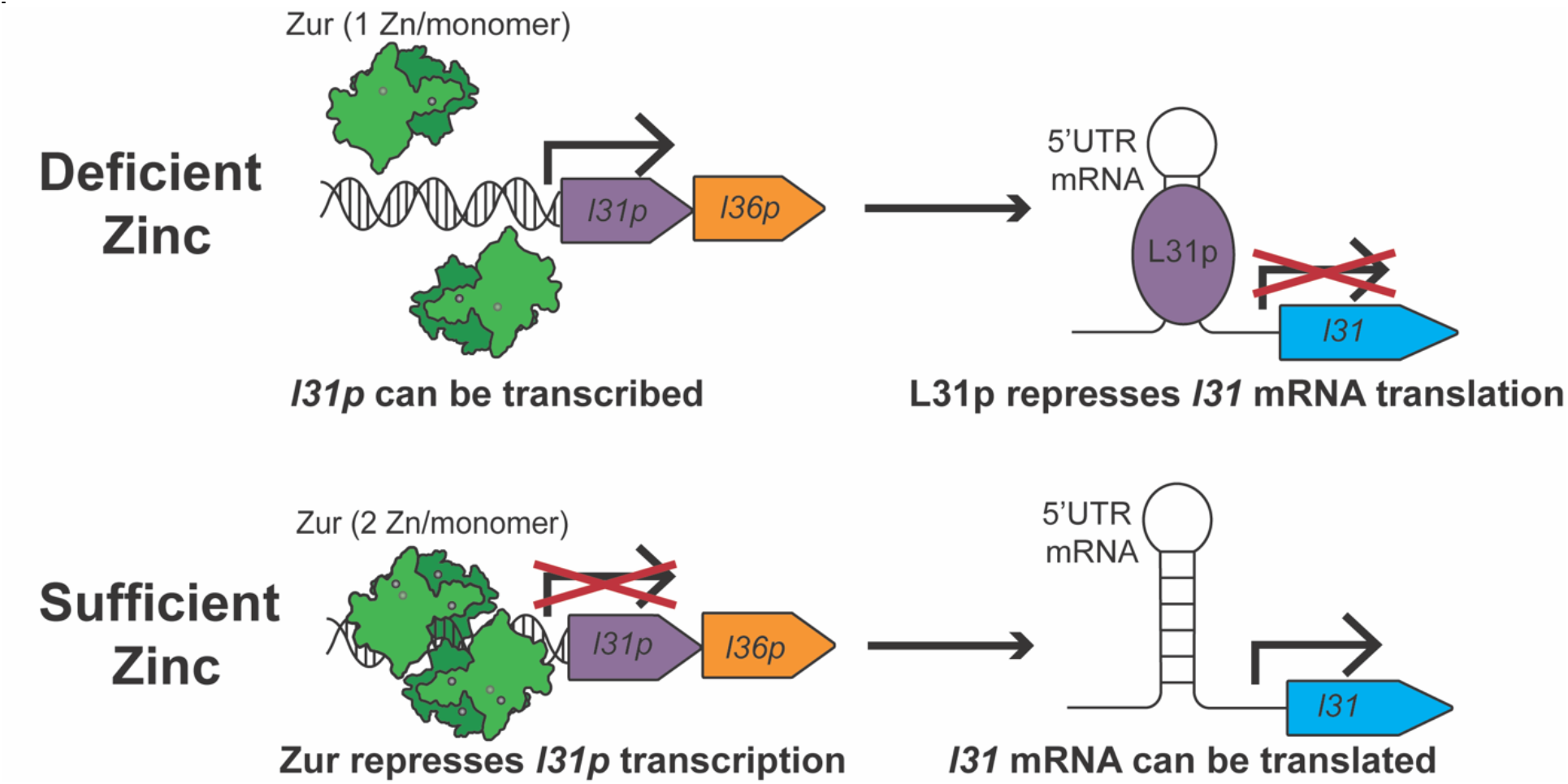
Schematic for proposed mechanism for Zur and zinc regulating L31 protein expression levels through a network involving L31p. When free Zn^2+^ is below subfemtomolar concentration, only one zinc is bound to each monomer of Zur, allowing L31p to be expressed. We propose that L31p represses L31 translation by binding to a 5’ UTR structure in the *l31* mRNA. At higher free zinc levels, Zur is bound to zinc ions, causing it to repress L31p expression. Depleting L31p thus allows L31 translation. Overall, we propose that L31p acts to invert the action of Zur by converting Zn-mediated repression into a genetic activator of L31 expression.

This information alone is not enough to determine if the repression of L31 by L31p occurs through transcription, translation, or RNA-degradation-level regulation. RNA-binding proteins can modulate any of these levels or multiple at once (52). Even though we observe a decreased *L31-sfGFP* mRNA level in *Δzur* cells (Figure 3A), translational regulation of the transcript could still occur. Reduced translation can increase a message’s susceptibility to RNAse degradation or increase transcription termination, as ribosomes and RNA polymerase can occupy the same message simultaneously (47–50) Several instances of autoregulatory ribosomal protein that primarily act through translational mechanisms also decrease its mRNA half-life in *E. coli*, including L1 and S4 (57, 58). *E. coli* ribosomal protein L4 can regulate its own expression at both at the translational level and by regulating the formation of a transcription terminator in its operon (59, 60). Further studies are needed to determine at which of these levels L31p regulates *l31*.

Mutational analyses of the *l31* 5’UTR and L31p protein provide several key insights to the proposed L31p protein-*l31* mRNA binding mechanism. The longest stem-loop in the *l31* 5’UTR appears to be most important for this regulation, which parallels the findings of L31 autoregulation (Figure 6C-D) (27). Deleting nucleotides 74 and 76 from bulge 2 in that stem-loop broke the L31-sfGFP on at a high level, while just mutating nucleotides 74-76 resulted in L31-sfGFP regulation more similar trend--although not identical--to the native *l31* 5’UTR. (Figure 6F). Therefore, maintenance of the overall structure of the bulge appears to be more important than specific nucleotide identity. Based on the NUPACK prediction, two base pairs between the two bulges have a lower predicted probability of pairing compared to other base pairs within the stem-loop (Figure S6). This could result in a dynamic opening of those base pairs to create one large bulge from the two smaller bulges. In a U54C mutant, which increases the probability of base pairing by replacing a U-G wobble pair with a C-G Watson-Crick-Franklin pair, the percent change to *Δzur* was almost completely eliminated (Figure 6F). Wobble U-G base pairs serve many necessary roles in RNA structures due to their conformational flexibility and alteration of the helical twist compared to canonical pairs (61). Similar asymmetric bulges adjacent to non-canonical base pairs are present in other bacterial ribosomal protein mRNA that bind ribosomal proteins, including the *E. coli* L1 and L10(L12)_4_ binding sites (28, 62, 63). Combined, this information could indicate that L31p protein binds to the *l31* 5’UTR at this region at bulge 2 when it is correctly oriented, and it depends on the increased flexibility from being adjacent to a G-U wobble pair.

Mutational and biochemical analyses of the *l31* 5’UTR and L31p protein provides limited insights into *l31* mRNA-L31p interaction. For one, our study and previous analyses of the *l31* 5’UTR involves the predicted minimum free energy equilibrium structure. While this structure can play an important role in this mechanism, other RNA structures, including higher energy structures in equilibrium and co-transcriptional RNA folding pathways, can also be critical to regulatory mechanisms (64, 65). Consequently, more rigorous structural studies are needed to determine more detailed structural, dynamic, and mechanistic information.

Results of the *in vivo* assays suggest that L31p appears to be the primary repressor of L31-sfGFP. Still, we cannot rule out the possibility that L36p, which is encoded in the same operon as L31p, also plays a role in this mechanism. L36p could contribute to L31p’s repression of L31; mechanisms of two ribosomal proteins binding to one transcript have been reported (66, 67). The S6:18 and L10(L12)_4_ proteins are proposed to regulate translation of their transcripts by binding to their mRNA, indicating this as possible for L31p and L36p (66, 67). *In vitro* studies with both L31p and L36p proteins are needed to determine if L36p could play an additive role in this mechanism.

Our L31/L31p switching mechanism in Figure 8 complements some recent studies that elucidate the function of zinc-lacking paralogues in bacterial ribosomal proteins. Recent work has shown that L31p (or L36p) in the ribosome allows improved growth and translation compared to cells lacking either both L31 and L31p or L36 and L36p (13). Compared with L31, though, L31p moderately reduces low-temperature growth, translational fidelity within the reading frame, and translation processivity in cells with L31p (12). A study in *Mycobacteria smegmatis* used ribosome profiling to determine that the zinc-lacking ribosomal proteins preferentially translate certain genes compared to the zinc-binding forms (18); however, *Mycobacteria smegmatis* have a different set of C+/C- paralogue ribosomal protein compared to *E. coli* (8). Profiling ribosomes in *E. coli* with L31/L36 or L31p/L36p could provide insight on the role of L31p in translation specificity. These and other methods to analyze ribosome function could help explain the significance of bacteria switching to these paralogues in zinc-deficient conditions.

In conclusion, the transcription factor Zur and its ligand zinc increase expression of zinc-dependent ribosomal L31 protein in *E. coli* in a multistep process. The data support a model in which regulation of L31 protein occurs through a repression of a repressor mechanism: Zur and zinc repress transcription of ribosomal protein gene *l31p*, and L31p protein represses translation of L31 protein expression by binding to the *l31* mRNA 5’UTR. This mechanism helps to explain how bacteria can change ribosome composition to adapt to environmental stress.

## Data Availability

Data generated in this manuscript are available in the extended data file.

## Supplementary Data

Supplementary data are available in the supplemental information.

## Acknowledgements

We thank Steven Philips for assisting with ribosome purification and Suning Wang for the unpublished results that inspired the initiation of this project. Elemental analysis was performed at the Northwestern University Quantitative Bio-element Imaging Center generously supported by NASA Ames Research Center Grant NNA04CC36G. Proteomic analysis was performed at the Northwestern Proteomics Core Facility, supported by NCI CCSG P30CA060553 awarded to the Robert H Lurie Comprehensive Cancer Center, instrumentation award (S10OD025194) from NIH Office of Director, and the National Resource for Translational and Developmental Proteomics supported by P41GM108569. Molecular graphics and analyses performed with UCSF ChimeraX, developed by the Resource for Biocomputing, Visualization, and Informatics at the University of California, San Francisco, with support from National Institutes of Health R01-GM129325 and the Office of Cyber Infrastructure and Computational Biology, National Institute of Allergy and Infectious Diseases.

