## Supplementary Information for "Zur and Zinc Increase Expression of *E. coli* Ribosomal Protein L31 Through RNA-Mediated Repression of the Repressor L31p"

[1]Interdisciplinary Biological Sciences Graduate Program, Northwestern University, Evanston, IL 60208, USA
[2]Chemistry of Life Process Institute, Northwestern University, Evanston, Illinois, 60208, United States of America.
[3]Northwestern Proteomics Core, Northwestern University, Evanston, Illinois, 60208, United States of America.
[4]Mass Spectrometry Technology Access Center, Washington University in St Louis, School of Medicine
[5]Department of Chemical and Biological Engineering, Northwestern University, Evanston, IL 60208, USA
[6]Center for Synthetic Biology, Northwestern University, Evanston, IL 60208, USA
[7]Department of Chemistry, Northwestern University, Evanston, Illinois, 60208, United States of America.
[8]Department of Chemistry, Michigan State University, East Lansing, MI 48824, USA.
[9]Department of Microbiology & Molecular Genetics, Michigan State University, East Lansing, MI 48824, USA.
*Corresponding authors

Table of Contents


| Figure | Page |
|---|---|
| **Figure S1.** ***In vivo*** **GFP assay growth calibration in WT and** ***Δzur E. coli*** **cells grown in LB.** | 2 |
| **Figure S2. Second replicate experiment of reverse-transcription quantitative PCR.** | 3 |
| **Figure S3. PAGE gels of monolith chromatography fractions of purified ribosomes from A.) WT and B.)** ***Δzur E. coli*** **cells.** | 3 |
| **Figure S4.** ***In vivo*** **reporter gene assays for fluorescence and growth of strains with** ***l31p*** **or** ***l36p*** **knocked out.** | 4 |
| **Figure S5. Growth of strains in zinc-deficient and zinc sufficient conditions.** | 5 |
| **Figure S6. Predicted secondary structures of the** ***l31*** **5'UTR and its mutants.** | 6-7 |
| **Figure S7. The top bulge of the** ***l31*** **5'UTR stem loop is not important for the regulation of L31-sfGFP by the** ***zur*** **gene.** | 8 |
| **Figure S8.** ***In vivo*** **sfGFP gene assays including plasmids that constitutively overexpress ribosomal proteins.** | 9 |
| **Table S1. Plasmids used in this study.** | 10 |
| **Table 2. ICP-MS measurements of Zn in LB media used in zinc-depletion experiments.** | 11 |
| **Table S3. Primers used in RT-qPCR.** | 11 |

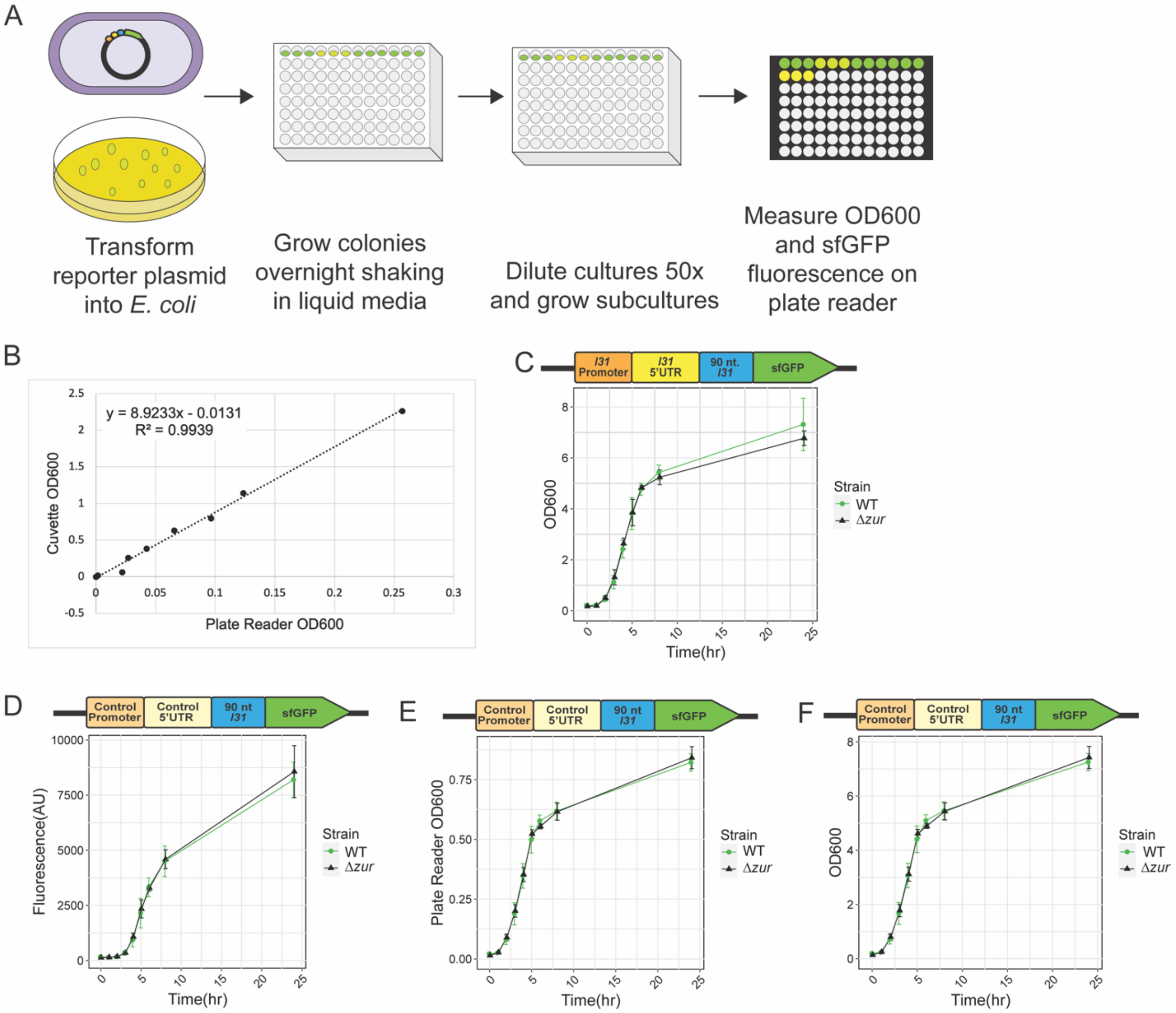


**Figure S1. *In vivo* GFP assay growth calibration in WT and *Δzur E. coli* cells grown in LB.** A.) Schematic for experiment setup of live cell reporter gene assays. B.) Calibration curve generated by diluting a saturated WT *E. coli* culture in LB and measuring both on the plate reader and a cuvette in a spectrophotometer. The linear regression calculation was generated by Microsoft Excel. C.) OD600 of cells with L31-sfGFP plasmid over time, using the equation from panel A to convert the plate reader measurements into the more standard spectrophotometer values. D.) Fluorescence from Control-sfGFP plasmid in cells from 0-24 hours, measured on a plate reader. E.) OD600 of cells with L31-sfGFP plasmid over time measured on the plate reader, and F.) converted to standard growth values using the equation from panel B. The bars indicate averages of three biological replicates (independent experiments), each performed with three technical replicates (cultures per experiment) for a total of nine data points (n=9.) The error bars represent standard deviation of the mean.

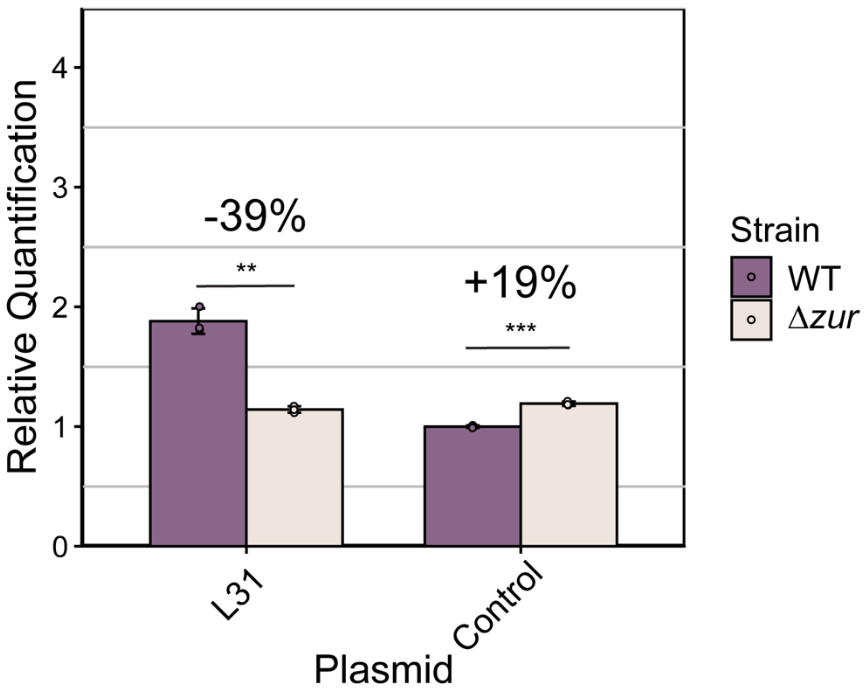


**Figure S2. Second replicate experiment of reverse-transcription quantitative PCR.**
See Figure 3A for details.

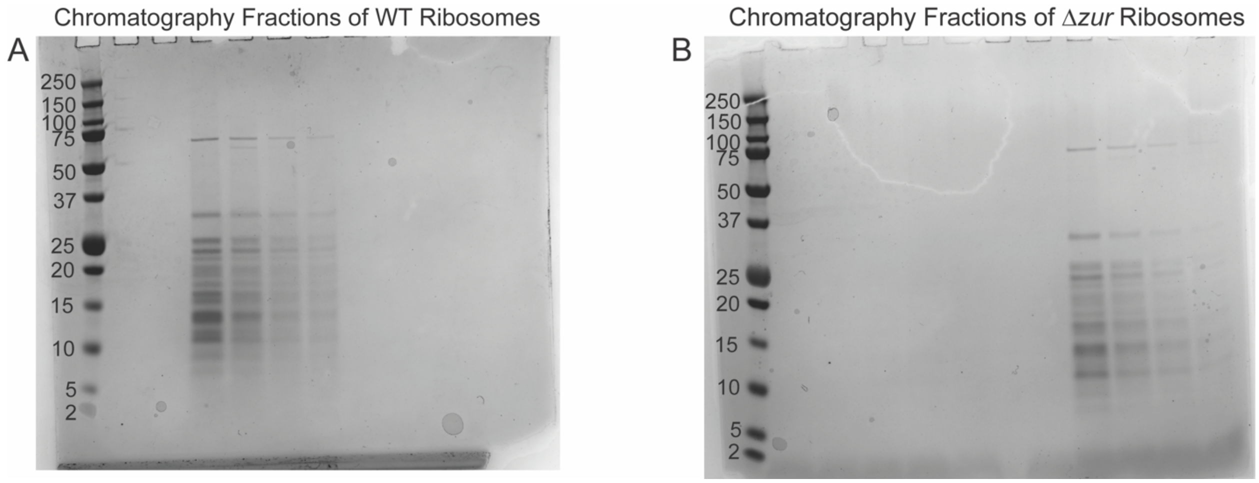


**Figure S3. PAGE gels of monolith chromatography fractions of purified ribosomes from A.) WT and B.) *Δzur E. coli* cells.**

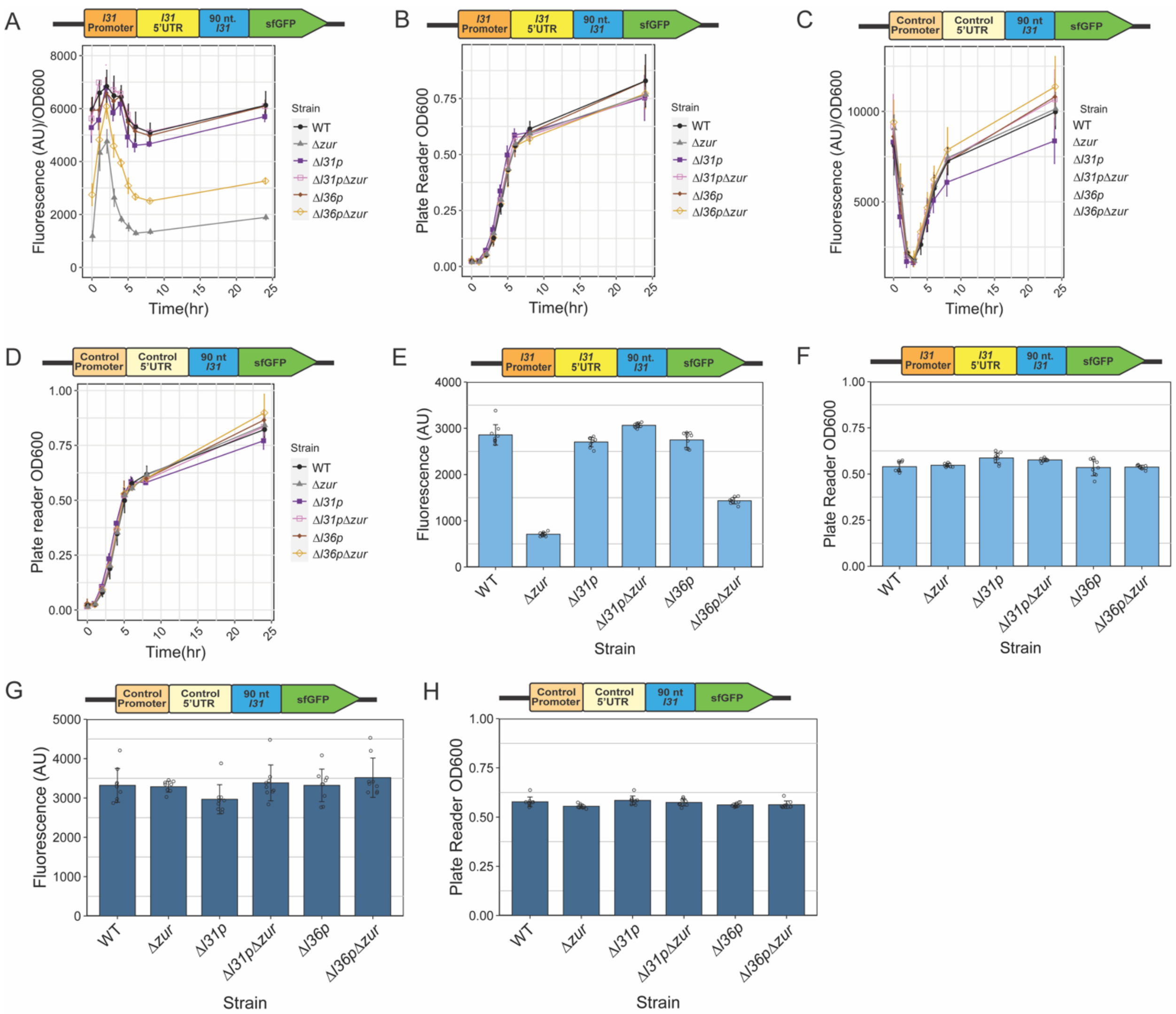


**Figure S4. *In vivo* reporter gene assays for fluorescence and growth of strains with *I31p* or *I36p* knocked out.** A.) Fluorescence/$OD_{600}$ and B.) plate reader $OD_{600}$ time-course of *E. coli* knockout strains with L31-sfGFP plasmid grown in LB media. C.) Fluorescence/$OD_{600}$ and D.) plate reader $OD_{600}$ time-course of *E. coli* knockout strains with control-sfGFP plasmid grown in LB media. E.) Plate reader fluorescence and F.) $OD_{600}$ of strains grown in LB with L31-sfGFP plasmid for 6 hr. G.) Plate reader fluorescence and H.) $OD_{600}$ of strains grown in LB with control-sfGFP plasmid for 6 hr. The bars indicate averages of three biological replicates (independent experiments), each performed with three technical replicates (cultures per experiment) for a total of nine data points (n=9.) The error bars represent standard deviation of the mean.

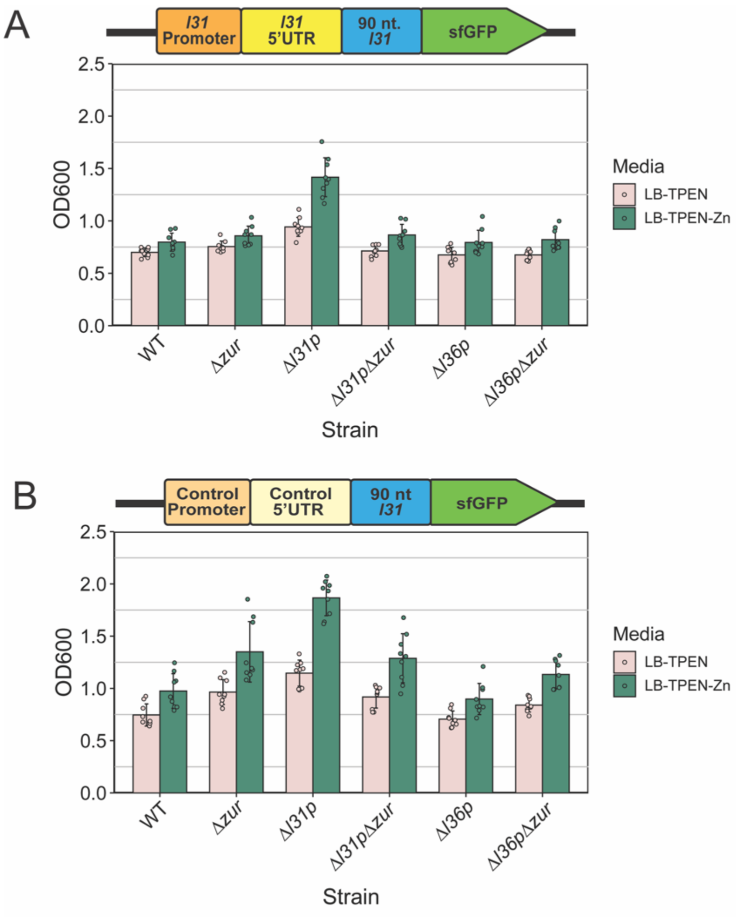


**Figure S5. Growth of strains in zinc-deficient and zinc sufficient conditions.** A.) Calibrated $OD_{600}$ of various strains with sfGFP plasmid, grown in LB + 100 μM of TPEN for 2 hours, then with or without the addition of 100 μM $ZnSO_4$ for an additional 2 hours. B.) Calibrated $OD_{600}$ of various strains with control sfGFP-plasmid, grown in LB + 100 μM of TPEN for 2 hours, then with or without the addition of 100 μM $ZnSO_4$ for an additional 2 hours. In each graph, the points indicate averages of three biological replicates (independent experiments), each performed with three technical replicates (cultures per experiment) for a total of nine data points (n=9.) The error bars represent standard deviation of the mean.

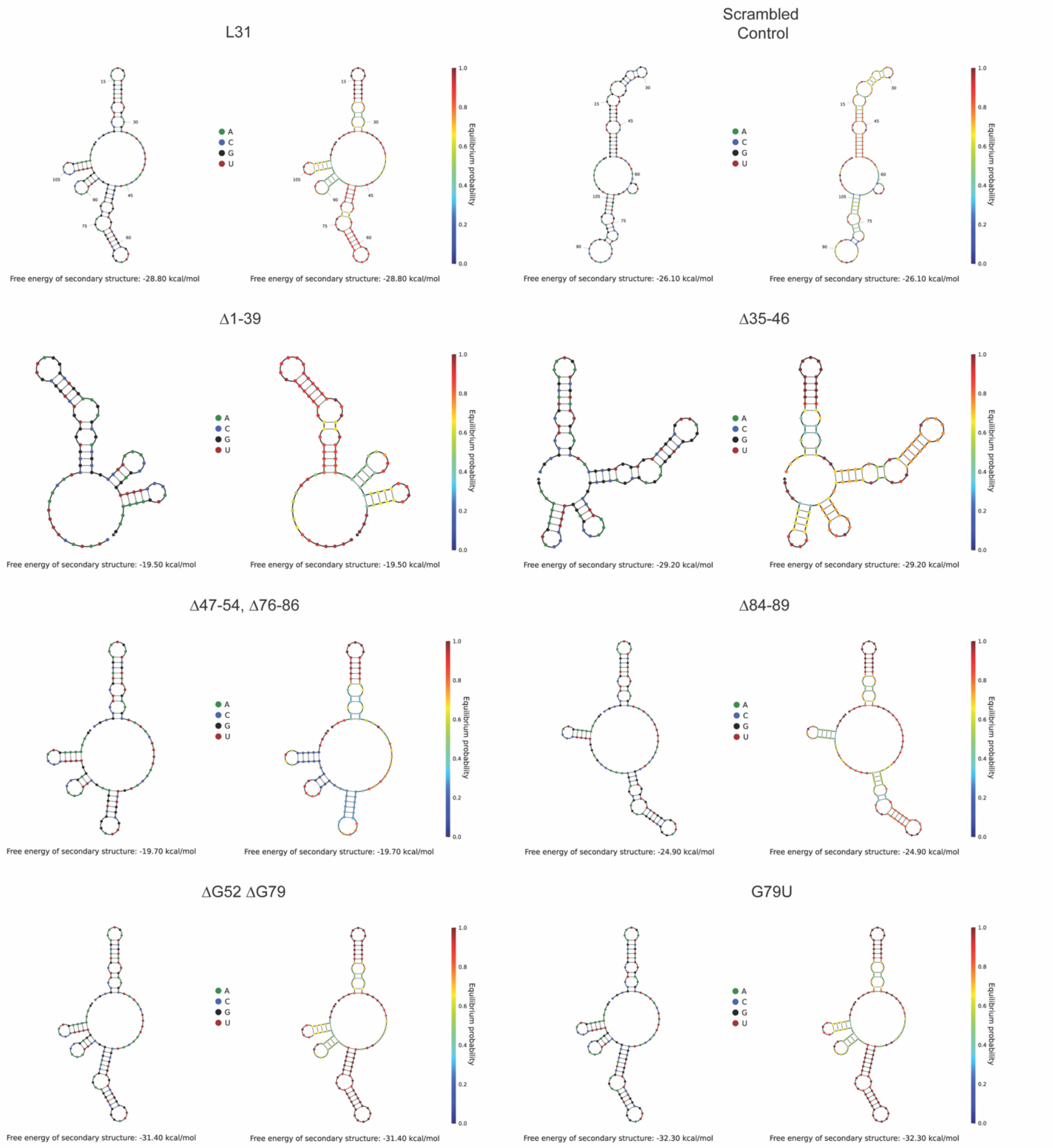
L31
Scrambled Control
Δ1-39
Δ35-46
Δ47-54, Δ76-86
Δ84-89
ΔG52 ΔG79
G79U
A
C
G
U
Equilibrium probability
Free energy of secondary structure: -28.80 kcal/mol
Free energy of secondary structure: -26.10 kcal/mol
Free energy of secondary structure: -19.50 kcal/mol
Free energy of secondary structure: -29.20 kcal/mol
Free energy of secondary structure: -19.70 kcal/mol
Free energy of secondary structure: -24.90 kcal/mol
Free energy of secondary structure: -31.40 kcal/mol
Free energy of secondary structure: -32.30 kcal/mol

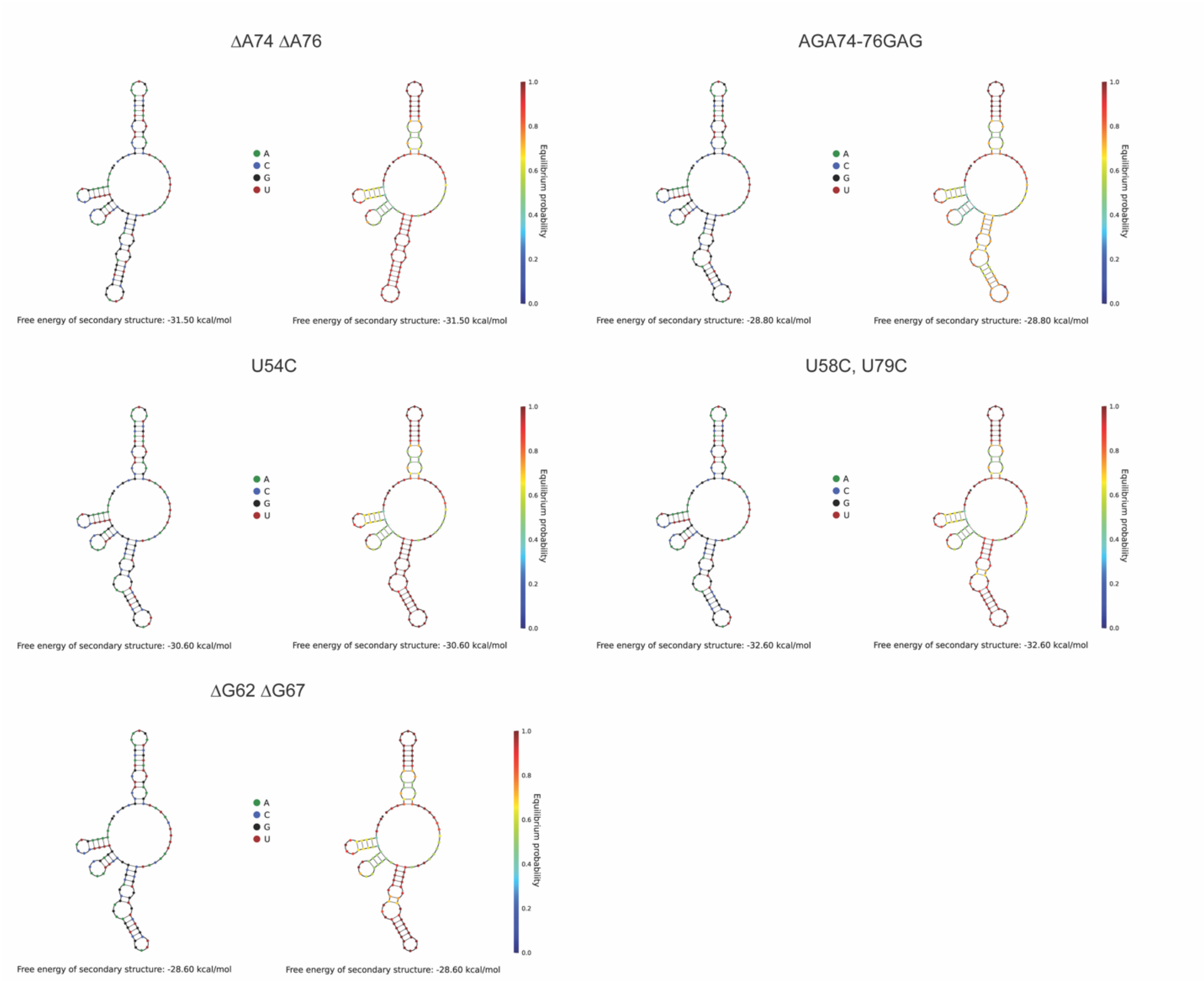


**Figure S6. Predicted secondary structures of the *I31* 5'UTR and its mutants.** Structures were generated using the webserver NUPACK, showing both sequence identity (left for each) and secondary structure probability (right for each). Parameters were set at temperature = 37 °C, number of strand species = 1, and maximum complex size = 1 strand.

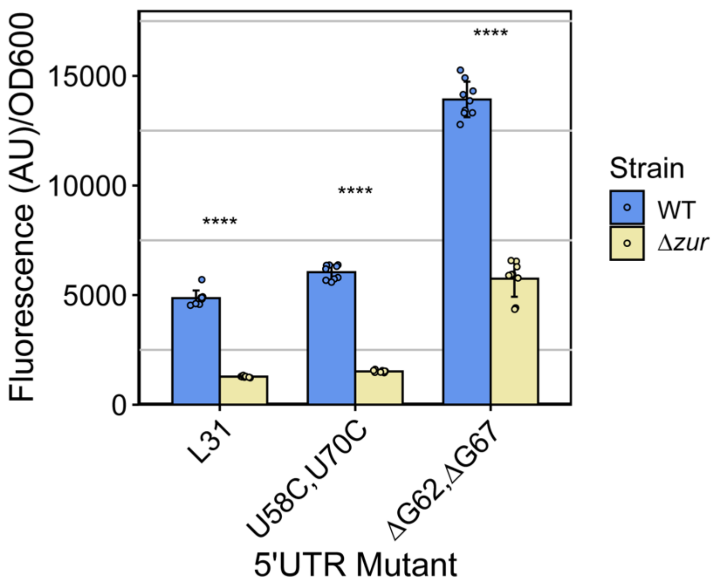


**Figure S7. The top bulge of the *l31* 5'UTR stem loop is not important for the regulation of L31-sfGFP by the *zur* gene.** Fluorescence/OD600 of cells grown in LB with *l31* 5'UTR mutations in L31-sfGFP plasmids. The bars indicate averages of three biological replicates (independent experiments), each performed with three technical replicates (cultures per experiment) for a total of nine data points (n=9.) The error bars represent standard deviation of the mean. Significance was calculated with a 2-tailed student's t-test between the fluorescence/$OD_{600}$ values for no added zinc vs added zinc for each strain. p-value < 0.05 = *, p-value < 0.01 = **, p-value < 0.001 = ***, p-value < 0.0001 = ****.

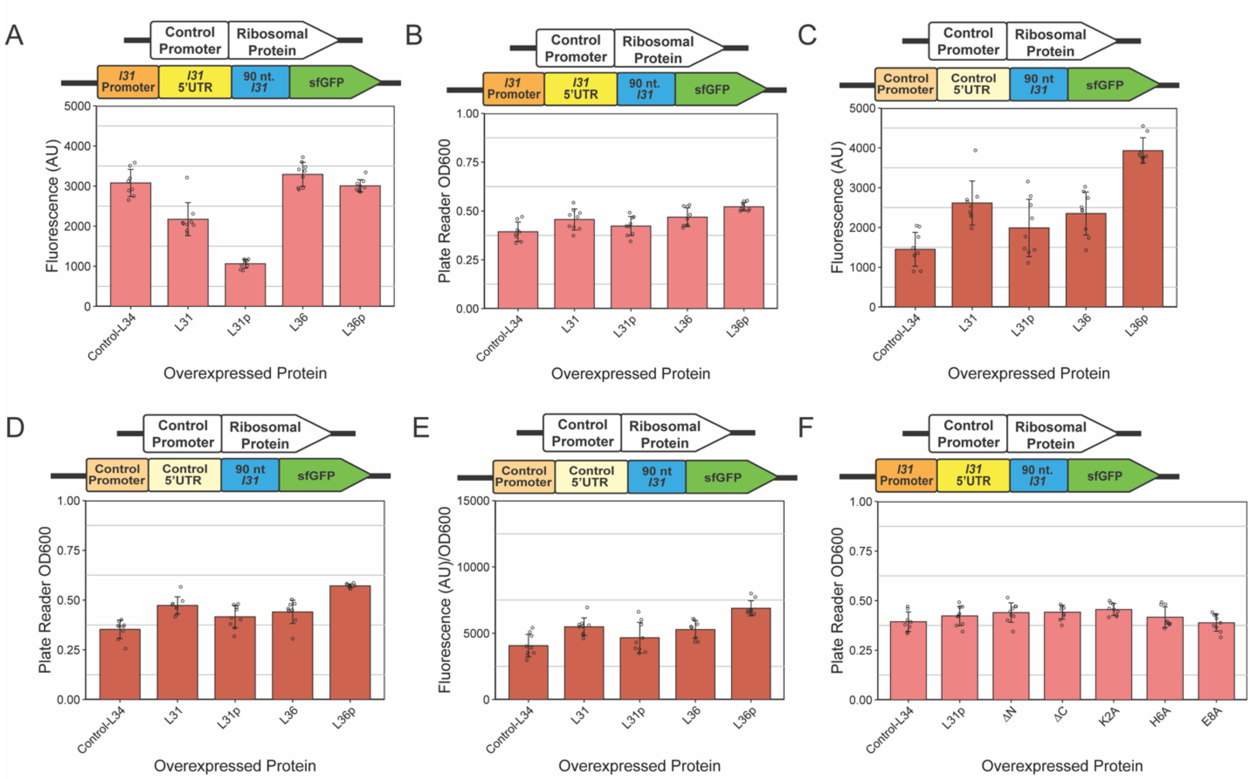


**Figure S8. *In vivo* sfGFP gene assays including plasmids that constitutively overexpress ribosomal proteins.** Cells were grown in LB for 6 hours. A.) Fluorescence (AU) and B.) OD600 measured on a plate reader for WT *E. coil* overexpressing various ribosomal proteins alongside the L31-sfGFP reporter plasmid. C-D.) Same as above but replacing the L31-sfGFP reporter plasmid with the control reporter plasmid and E.) Fluorescence/OD600with the control-sfGFP reporter plasmid. F.) Same a B.) but overexpressing L31p protein mutants. The bars indicate averages of three biological replicates (independent experiments), each performed with three technical replicates (cultures per experiment) for a total of nine data points (n=9). The error bars represent standard deviation of the mean.

**Table S1. Plasmids used in this study.**

| Plasmid | Function |
|---|---|
| pRAR017 | Reporter L31-sfGFP plasmid |
| pRAR039 | Reporter L31-sfGFP plasmid with nucleotides (nt) 1-31 of *l31* 5'UTR deleted |
| pRAR040 | Reporter L31-sfGFP plasmid with nt 35-46 of *l31* 5'UTR deleted |
| pRAR041 | Reporter L31-sfGFP plasmid with nt 47-54 & 76-86 of *l31* 5'UTR deleted |
| pRAR042 | Reporter L31-sfGFP plasmid with nt 84-89 of *l31* 5'UTR deleted |
| pRAR051 | Reporter L31-sfGFP plasmid with nt 47-51 & 80-84 of l31 5'UTR mutated |
| pRAR052 | Reporter L31-sfGFP plasmid with nt 56-61 & 68-73 of *l31* 5'UTR mutated |
| pRAR064 | Reporter L31-sfGFP plasmid with 5'UTR replaced with scrambled control |
| pRAR074 | Reporter L31-sfGFP plasmid with promoter replaced with constitutive J23108 and 5'UTR replaced with scrambled control |
| pRAR089 | *In vivo* expression of L36p with constitutive promoter J23108 |
| pRAR090 | *In vivo* expression of L36 with constitutive promoter J23108 |
| pRAR091 | *In vivo* expression of L31p with constitutive promoter J23108 |
| pRAR092 | *In vivo* expression of L31 with constitutive promoter J23108 |
| pRAR101 | *In vivo* expression of L31p with residues 2-8 deleted, constitutive promoter J23108 |
| pRAR102 | Reporter L31-sfGFP plasmid with U54C in *l31* 5'UTR |
| pRAR104 | Reporter L31-sfGFP plasmid with A74 and A76 in *l31* 5'UTR |
| pRAR105 | Reporter L31-sfGFP plasmid with G52 and G79 deleted in *l31* 5'UTR |
| pRAR106 | Reporter L31-sfGFP plasmid with G79U in *l31* 5'UTR |
| pRAR107 | Reporter L31-sfGFP plasmid with AGA 74-76 GAG mutation in *l31* 5'UTR |
| pRAR109 | Reporter L31-sfGFP plasmid with G60 and C69 deleted in *l31* 5'UTR |
| pRAR110 | Reporter L31-sfGFP plasmid with G62 and G67 deleted in *l31* 5'UTR |
| pRAR111 | *In vivo* expression of L31p with residues last 8 C-terminal residues deleted, constitutive promoter J23108 |
| pRAR112 | *In vivo* expression of L31p with K2A mutation, constitutive promoter J23108 |
| pRAR113 | *In vivo* expression of L31p with H6A mutation, constitutive promoter J23108 |
| pRAR114 | *In vivo* expression of L31p with E8A mutation, constitutive promoter J23108 |
| pRAR116 | Reporter L31-sfGFP plasmid with U58C and U70C in *l31* 5'UTR |
| pRAR119 | *In vivo* expression of L34 with constitutive promoter J23108 |
| pRAR120 | Reporter L31-sfGFP plasmid with promoter replaced J23108 |

**Table 2. ICP-MS measurements of Zn in LB media used in zinc-depletion experiments.**

| ICP-MS LB sample | Calculated Zn Concentration (μM) |
|---|---|
| **1** | **92.3** |
| **2** | **91.0** |
| **3** | **91.2** |
| **Average** | **91.5** |

**Table S3. Primers used in RT-qPCR.**

| Primer | Sequence |
|---|---|
| sfGFP reverse transcription | TTATTTGTAGAGCTCATCCATG |
| 16S rRNA reverse transcription | TAAGGAGGTGATCCAACCG |
| sfGFP qPCR forward | CACTGGAGTTGTCCCAATTCT |
| sfGFP qPCR reverse | TCCGTTTGTAGCATCACCTTC |
| 16S rRNA qPCR forward | GTCAGCTCGTGTTGTGAAATG |
| 16S rRNA qPCR reverse | CCCACCTTCCTCCAGTTTATC |
